# A minimal physical model of adhesion-dependent protrusion explains confined haptotaxis and oscillatory cell migration

**DOI:** 10.64898/2026.07.30.741711

**Authors:** Jeevan Kumar Papaganti, Jonathan E. Ron, Shubhadeep Sadhukhan, Isabela Corina Fortunato, Steffen Grosser, Ushasi Roy, Raimon Sunyer, Xavier Trepat, Nir S. Gov

## Abstract

Haptotaxis is the directed migration of cells along gradients of substrate adhesion, and is an important guidance mechanism in biology. Recent experiments on fibronectin-patterned tracks show that cells typically migrate toward higher adhesion, then oscillate around the adhesion maximum, with substantial variability in speed and cell length. To explain these cellular shape and migration dynamics we present a minimal model for confined haptotaxis, built around one key physical ingredient: local adhesion strength directly enhances the recruitment of protrusive actin polymerization activity at the cell edge. This coupling creates a front–rear difference in protrusive activity, biasing the cell towards polarization in the direction of the higher adhesion. Quantitative comparison with experiment shows strong agreement at multiple levels: cell population-level directionality statistics and position-dependent changes in cell length and velocity. At the single-cell level, the model reproduces the full diversity of experimentally observed trajectories, including haptotactic bias and its dependence on adhesion-gradient strength, initial position, length and speed variations, myosin II inhibition, and migration on inverted adhesion gradients. Variability in experimental trajectories and migration speed is explained by differences in intrinsic actin polymerization activity and stochastic fluctuations, accounting for the full spectrum of observed migration patterns. Together, our results identify adhesion-dependent amplification of protrusive activity as a minimal and sufficient physical mechanism for understanding haptotaxis.

## I. INTRODUCTION

Directed cell migration is central to embryonic morphogenesis, immune responses, wound healing, and cancer invasion [1–3]. Cells achieve directional motion by detecting and responding to spatial cues in their environment, including gradients of soluble chemoattractants and gradients of substrate-bound adhesion ligands [4, 5]. Migration guided by spatial variations in substrate adhesiveness is commonly referred to as *haptotaxis* [6–8]. A closely related guidance mechanism is durotaxis, in which cells migrate along gradients of substrate stiffness [9, 10]. Because adhesion strength can increase with substrate stiffness, stiffness gradients can effectively act as adhesion gradients [11]. However, the stiffness–adhesion relationship is complex and context-dependent [12], making it difficult to isolate the contribution of adhesion alone. This motivates the study of pure adhesion gradients, in which the spatial cue is implemented directly at the level of ligand density, decoupling adhesiveness from stiffness and allowing to study the gradient-sensing mechanisms at the cell–ECM interface [13].

Recently, Fortunato *et al*. introduced an experimental platform to probe confined haptotaxis using fibronectin-patterned one-dimensional tracks with prescribed adhesion profiles [14] (Fig.1a,b). In these experiments, most cells initially migrated up the adhesion gradient — a response termed “haptotactic first runs” — yet a substantial fraction later migrated down the gradient, and many trajectories displayed long-lived oscillations around the adhesion maximum [14]. Note that these oscillations are distinct from those that are dominated by cellular-driven modifications due to deposition of biochemical signals or mechanical remodelling the extracellular matrix [15].

**Figure 1.**
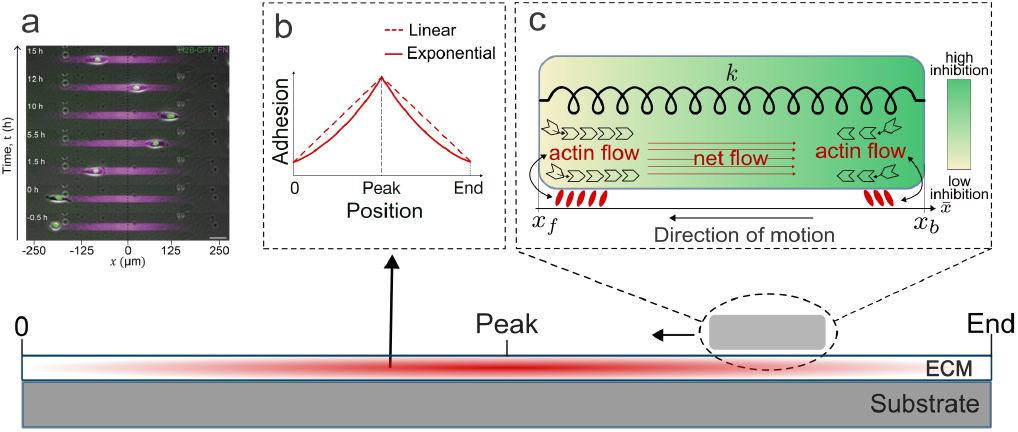
Cell migration on adhesion gradient tracks. The red color gradient (bottom illustration) represents adhesion strength on the extracellular matrix (ECM) coated on the substrate. **a**) Representative experimental kymograph of cell migration on an adhesion gradient over 15 hours. The fibronectin concentration gradient is represented by magenta colour intensity. Cell nucleus position (green) is plotted at different times. The cell track spans 500 *µ*m, centered at *x* = 0. **b**) Adhesion gradient profiles used in the theoretical model, as a function of position, showing linear (dashed line) and exponential (solid line) increase from the edges of the pattern to the peak at the center. **c**) Illustration of the model. Above, zoom in of the model mechanics. Below, a cell migration toward higher adhesion concentration (positive haptotaxis). Red arrows represent net actin retrograde flow. Actin-adhesion coupling is depicted by black arrows, with adhesions shown as red ovals and actin polymerization activity at the cell edges is indicated by black monomers, where their direction and number represent the local flow magnitude at the rear (*x*_*b*_) and front (*x*_*f*_). The green-to-yellow gradient represents spatial variation in the inhibitor concentration.

Beyond directional statistics, the experiments reported systematic position-dependent trends in cell shape and migration speed during up-versus down-gradient migration, pronounced cell-to-cell variability, and sensitivity to pharmacological perturbations of actomyosin contractility [14]. These observations raise fundamental questions: which minimal physical ingredients are sufficient to generate both directional bias and oscillatory migration on confined adhesion gradients, and how adhesion-dependent feedbacks interact with polarity, mechanics, and stochasticity to shape the observed cellular trajectories.

To address these questions, we present here a theoretical model, based on the coarse-grained one-dimensional framework of Ron *et al*. [16]. In this model a cell spontaneously polarizes through a feedback between actin retrograde flow that emanates from the opposing poles of the cell, and the advection and redistribution of a polarity cue [16, 17]. This was termed the Universal Coupling between Cell Speed and Persistence (UCSP) mechanism. This class of models accounts for persistent migration and can generate multiple motility modes — including stick-slip migration and motion with a fixed length — through the coupling between actin-driven protrusion, adhesion dynamics, and cell elasticity [16, 18]. Related one-dimensional frameworks based on active gel theory have also shown that mechanosensitive adhesion dynamics alone can drive symmetry breaking and haptotaxis on adhesion gradients [14, 19]. We aim to extend our theoretical understanding of haptotaxis using a model with explicit length and adhesion-polarity coupling, to describe the huge variability of cell migration patterns.

Our model goes beyond the theoretical description of haptotaxis given in [14], in several key aspects. In [14] the cell polarity was either fixed or purely affected by random noise, and therefore direction changes were only initiated by the cell reaching the limits of the adhesion range, or due to noise. Our model provides an explicit mechanism for self-polarization, namely the UCSP framework [16], thereby explaining the origin of the observed spontaneous polarity reversals and different oscillatory migration patterns observed in experiments on the symmetric adhesion pattern (Fig.1a). The complex migration patterns observed in experiments are explained naturally in our model as arising from the interplay between the polarity persistence and the response to the adhesion gradient. In addition, our model enables to naturally resolve smooth-stick-slip migration patterns and their transitions, as well as their traction force patterns, both not included in the model of [14].

Here, we extend the framework of [16] to describe the effects of spatially varying adhesion. In that framework, adhesion influences migration in two distinct ways: first, by setting the effective frictional resistance that couples cell motion to the substrate [16]; and second, by modulating the efficacy with which actin-driven protrusion is mechanically transmitted through the actin–adhesion ‘clutch’ to generate edge advancement [18]. The central new ingredient is an explicit coupling between local adhesion strength and the effective actin protrusive activity at each cell edge (Fig. 1c). This choice is motivated by molecular evidence that focal adhesions act as signaling and mechanical hubs that promote actin assembly and protrusive dynamics [8, 20–2]

In addition to biochemical signaling, adhesion can modulate actin assembly through a geometric mechanism at the cellular scale. Increased local adhesion promotes enhanced membrane spreading at the corresponding cell edge, effectively enlarging the polymerization-competent interface over which actin networks can assemble and push against the membrane. This coupling is bidirectional: elevated actin polymerization and the associated cytoskeletal organization promote the formation and maturation of adhesions, establishing a positive feedback between actin assembly and adhesion strengthening [24, 25].

This geometric effect of adhesion on the leading-edge recruitment of actin polymerization activity naturally arises in our recent three-dimensional mechanochemical simulations [26]. In this framework, a motile “minimal cell” emerges as a self-organized shape through the coupling of curved membrane complexes to actin-polymerization–driven protrusive forces, acting in the presence of substrate adhesion [27, 28]. The model additionally incorporates the UCSP feedback between the activity at the membrane and the concentration of a long-range inhibitory polarity cue that is advected by the global actin retrograde flow, thereby stabilizing persistent front–rear polarization via a feedback between flow-induced cue gradients and local protrusive activity [26].

Within this three-dimensional setting, imposing an adhesion energy gradient (parameterized by an increment Δ) produces a pronounced front–rear asymmetry in morphology and dynamics when combined with sufficiently strong advection of the inhibitor polarity cue (parameterized by *β*, Eq.1). The simulations (Fig.2) exhibit increased spreading area on the higher-adhesion side and a natural enhancement of protrusive activity biased toward the higher adhesion, relative to uniform adhesion or lack of internal advection activity. The simulated asymmetry compares well with experimental observations of migrating cells on adhesion gradients (Fig.2c), which similarly display a larger and more protrusive lamellipodium oriented up the fibronectin gradient. These results provide independent support for the central hypothesis that spatial variations in adhesion bias actin-driven protrusive activity, thereby locally amplifying the effective polymerization capacity.

**Figure 2.**
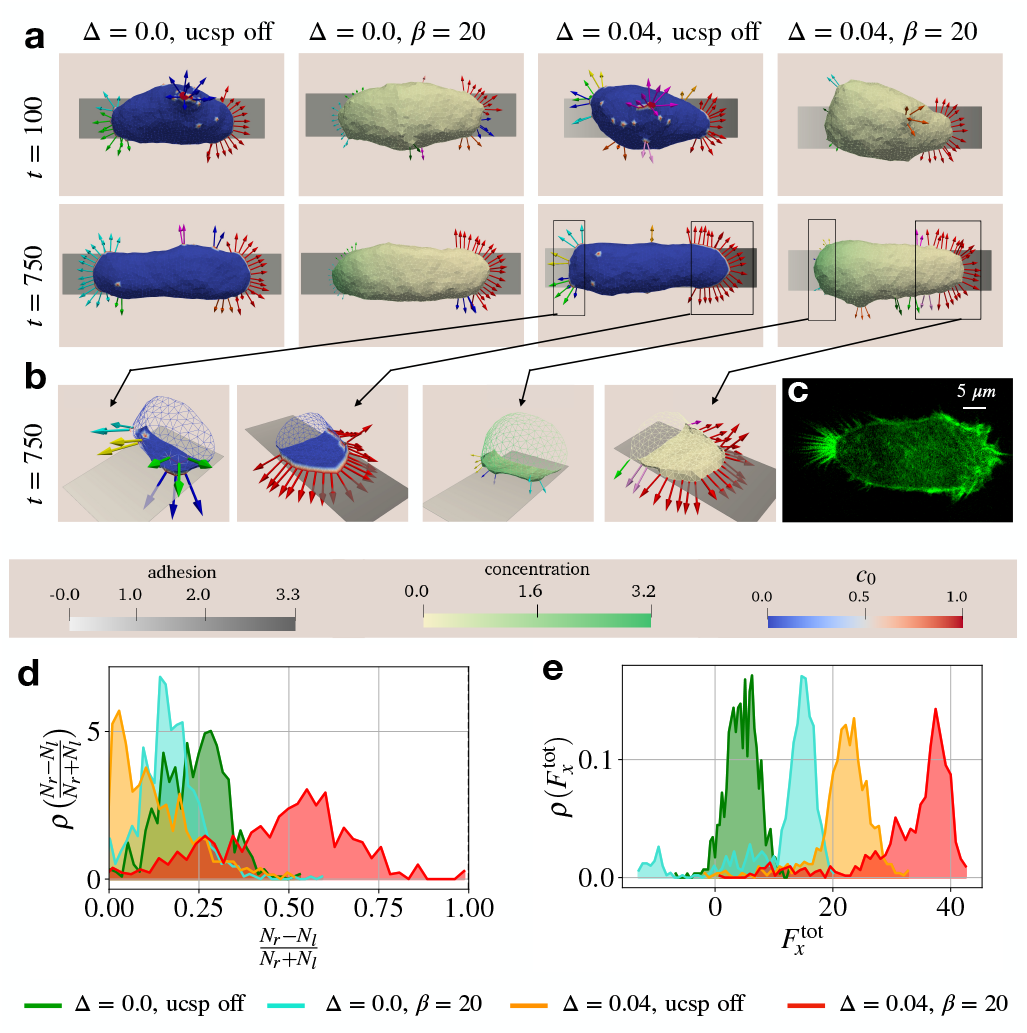
**a**) Snapshots of the simulated cell on the adhesive line (grey strip), with uniform adhesion (Δ = 0) or with a linear gradient 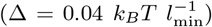 in adhesive strength. For each case, we show a simulation with and without the UCSP (self-polarization) mechanism. **b**) Enlarged view of the leading edge cluster formation at the rear and the front when the gradient in the adhesive patch is present. **c**) Representative experimental fluorescence image of an MCF10A cell expressing LifeAct-GFP (green) and migrating up the fibronectin gradient (increasing to the right), exhibiting a larger protrusive lamellipodia at the leading-edge facing the higher adhesion. **d**) Probability distribution function for the asymmetry in the number of curved proteins at the front and the back for the four different cases in (**a**). **d**) Probability distribution function for the total force, including the passive adhesive pull due to the adhesion gradient, for the cases in (**a**). We set the adhesive energy per node for the uniform case as *E*_ad_ = 1.5*k*_*B*_*T*. For the case of gradient in adhesion, we set the adhesive energy per node to be: *E*_ad_ = 0.5 + Δ(*x* − *x*_min_). We set the vesicle parameters as follows [26]: Number of nodes *N* = 1447, density of curved-active nodes *ρ* = 3.45%, active force magnitude 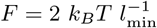, 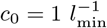, *c*_tot_ = 4000, *D* = 4000, *c*_*s*_ = 1.

We show here that the proposed minimal adhesion–actin coupling, integrated into the one-dimensional polarity-and-force-balance model, is sufficient to reproduce a broad set of experimental observations on exponential adhesion gradients [14]. These results provide a mechanistic account of confined haptotaxis and oscillatory migration, and offer a quantitative framework for exploring how adhesion-dependent feedbacks shape cell migration dynamics on patterned substrates.

## II. MODEL

### A. Actin-adhesion coupling

We incorporate actin–adhesion coupling into the one-dimensional framework [16, 29] by allowing the maximal actin polymerization speed at each cell edge to depend on the local adhesion strength (Fig. 1b,c). Denoting the two edges by subscripts *f* (front) and *b* (back), we write

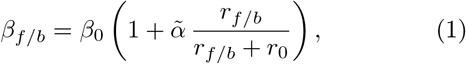

where *β*_0_ is the baseline polymerization speed (which we term “activity parameter”), 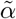 controls the enhancement factor of actin polymerization activity due to adhesion, and *r*_0_ is a characteristic adhesion scale that sets the onset of saturation. Varying *β*_0_ or 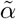 produces qualitatively similar effects on the migration dynamics, as both parameters primarily rescale the adhesion-dependent polymerization capacity. Therefore we fix 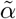 and use *β*_0_ as the principal control parameter for polymerization activity, which is responsible for the observed variability in cellular migration speeds.

### B. Equations of Motion

The protrusive force exerted by actin polymerization at the cell’s front depends on adhesion as

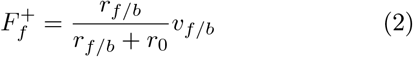

where *v*_*f/b*_ is the local actin treadmilling velocity, *r*_*f/b*_ is the adhesion affinity.

This protrusive force is opposed by a drag force and a restoring elastic force

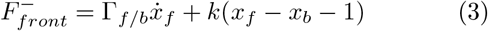

where Γ_*f/b*_ is the effective friction coefficient and *k* is the effective spring constant of the cell (including the effects of cell contractility), where the rest length is normalized to 1 (consistent with the rescaling of [16]).

Balancing these forces, both front and back, yields the position dynamics for the cell’s front and back

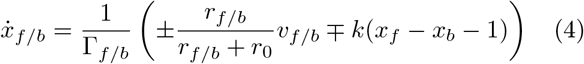

The friction coefficient, Γ_*f/b*_, depends on the direction of motion of the cell edge, whether it is protruding or retracting, and the dynamics of adhesion [18]

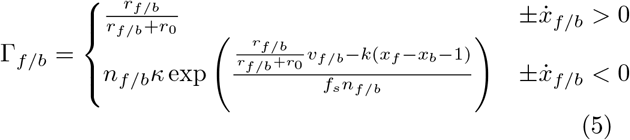

where *n*_*f/b*_ is the fraction of bound adhesion linkers, *κ* is the linker’s elastic stiffness, and *f*_*s*_ is the slip-bond force susceptibility. The fraction of bound adhesion linkers evolves according to

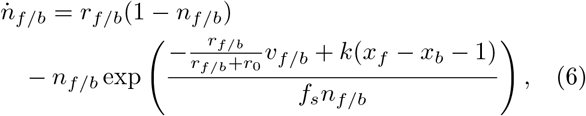

where the detachment rate depends on the force per linker (slip-bonds).

Finally, the local actin flow velocities relax toward their adhesion-dependent target values

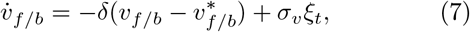

with

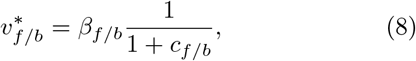

where *δ* is the relaxation rate and *σξ*_*t*_ refers to the uncorrelated Gaussian noise with an amplitude *σ*_*v*_ that is added to the actin flows. The concentration of the polarity cue at the cell edges is denoted by *c*_*f/b*_, which acts as an inhibitor of the actin polymerization activity.

The functional form of the inhibitor concentration at the cell edges *c*_*f/b*_ is given by the solution of the steady-state advection-diffusion equation [16, 17]. The values of this inhibitor at the two edges of the cell are given by

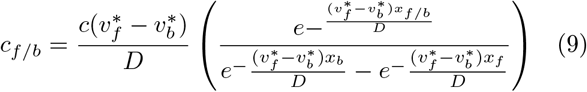

where *D* is the diffusion coefficient of the polarity cue and *c* is a dimensionless quantity given by the ratio of the total amount of polarity cue and its saturation concentration. This ties 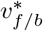 to *β*_*f/b*_, and thus to *r*_*f/b*_, completing the feedback mechanisms that determine the cell polarization coupling.

The model parameters used in this paper are summarized in Table I. We give there the parameters that do not change between cells, and were calibrated to fit the observed behavior of different migratory cells [16, 30, 31]. In our model cells spontaneously elongate when spreading on an adhesive surface. Above a critical value of the activity parameter *β*_0_ (Eq.1) such spreading cells spontaneously self-polarize, break symmetry and become motile. Throughout the paper we keep *β*_0_ above this critical value, to focus on the behavior of migratory cells.

**Table 1.**
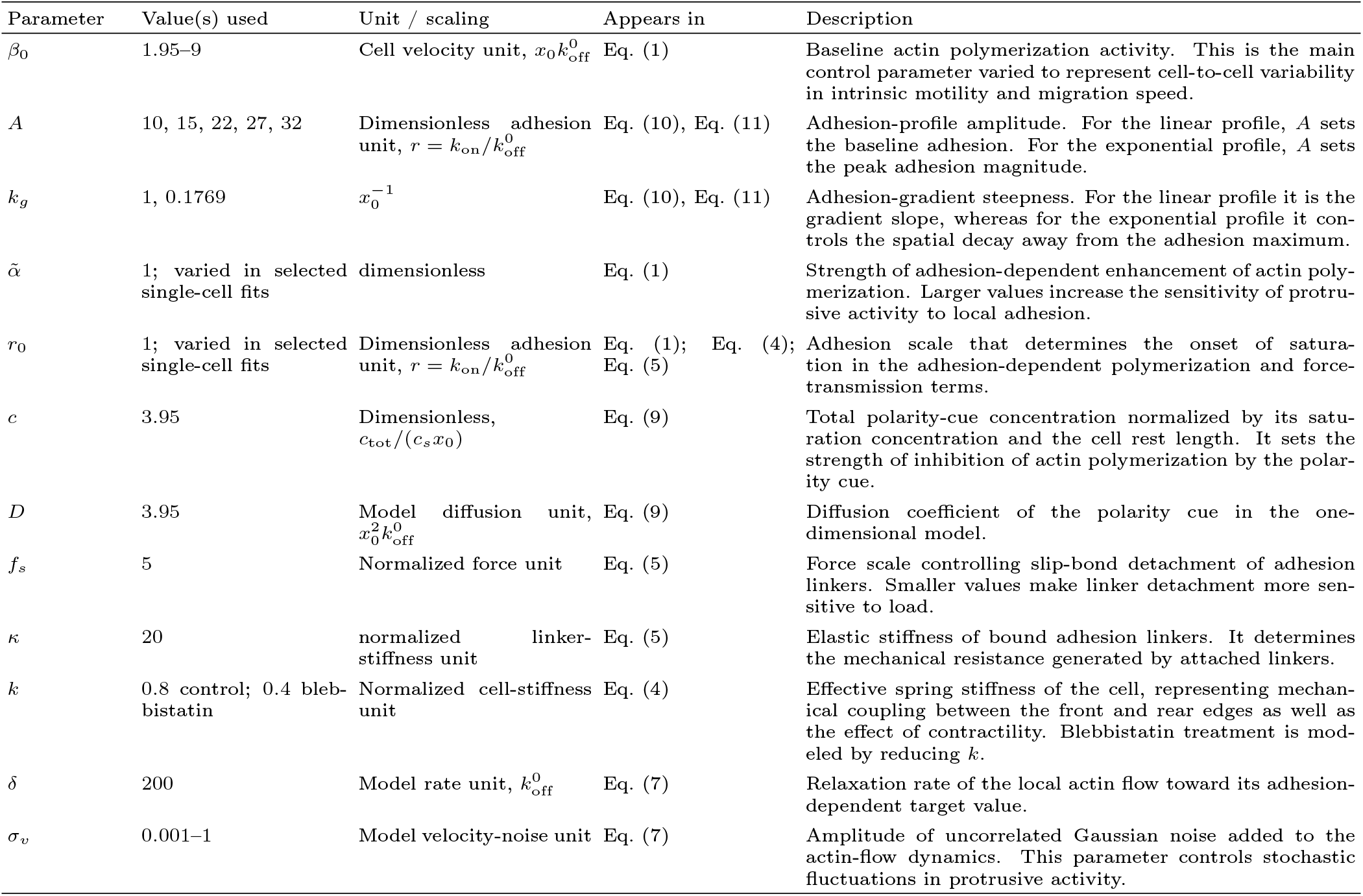
Model parameters used in the one-dimensional simulations. All values are reported in nondimensional model units following the scaling of Ron *et al*. [16]. The length scale in the problem is set by cell rest length *x*_0_ (which we set to be 1), the time scale is normalized by the bare rate of detachment of bound adhesion bonds 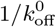, velocities are scaled by 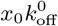, diffusion coefficients by 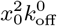, and adhesion is expressed as the dimensionless binding–unbinding ratio 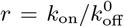. Unless otherwise stated, simulations on the exponential adhesion profile use: *A* = 27, *k*_*g*_ = 0.1769, *x*_peak_ = 0, 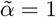, and *r*_0_ = 1.

The model length units are in terms of the rest length of the cell, before it spreads on the adhesive substrate. The model time units are normalized by the bare off-rate of the adhesion bonds (Eq.6). We set the parameters such that they cover the range of observed cellular migration and shape dynamics [16, 30, 31].

### C. Adhesion Gradient

We evaluated our model using two distinct adhesion gradient profiles: a linear profile and an exponential profile. The adhesion gradient profile exhibits a peak at *x* = *x*_peak_ and is symmetric in both directions (Fig. 1 **b**).

The linear adhesion gradient is:

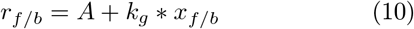

where *A* is the minimum adhesion and *k*_*g*_ is the gradient slope.

We introduce a symmetric double-exponential adhesion profile

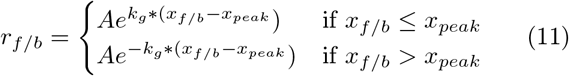

where *A* is the peak adhesion and *k*_*g*_ controls the decay rate. This symmetric double-exponential profile corresponds to the experimentally generated gradients [14] (Fig.1b).

## III. RESULTS

### A. Theoretical Predictions

#### 1. Linear adhesion gradient profile

We first examine cell migration on a linear adhesion gradient (Eq. (10), Fig.1b). The adhesion levels at the front and rear edges are denoted by *r*_*f*_ and *r*_*b*_, with Δ*r* ≡ *r*_*f*_ − *r*_*b*_ distinguishing up-gradient motion (Δ*r >* 0) from down-gradient motion (Δ*r <* 0) (Fig. 3**a**,**b**). Fig. 3c presents the phase diagram in the (*r*_*b*_, Δ*r*) parameter space. The red curve, computed by numerical continuation [32], marks the Hopf bifurcation separating smooth and stick–slip migration. At low baseline adhesion *r*_*b*_, stick–slip migration occurs even in the absence of an adhesion gradient, whereas increasing *r*_*b*_ requires larger adhesion gradients Δ*r* to sustain oscillatory migration.

**Figure 3.**
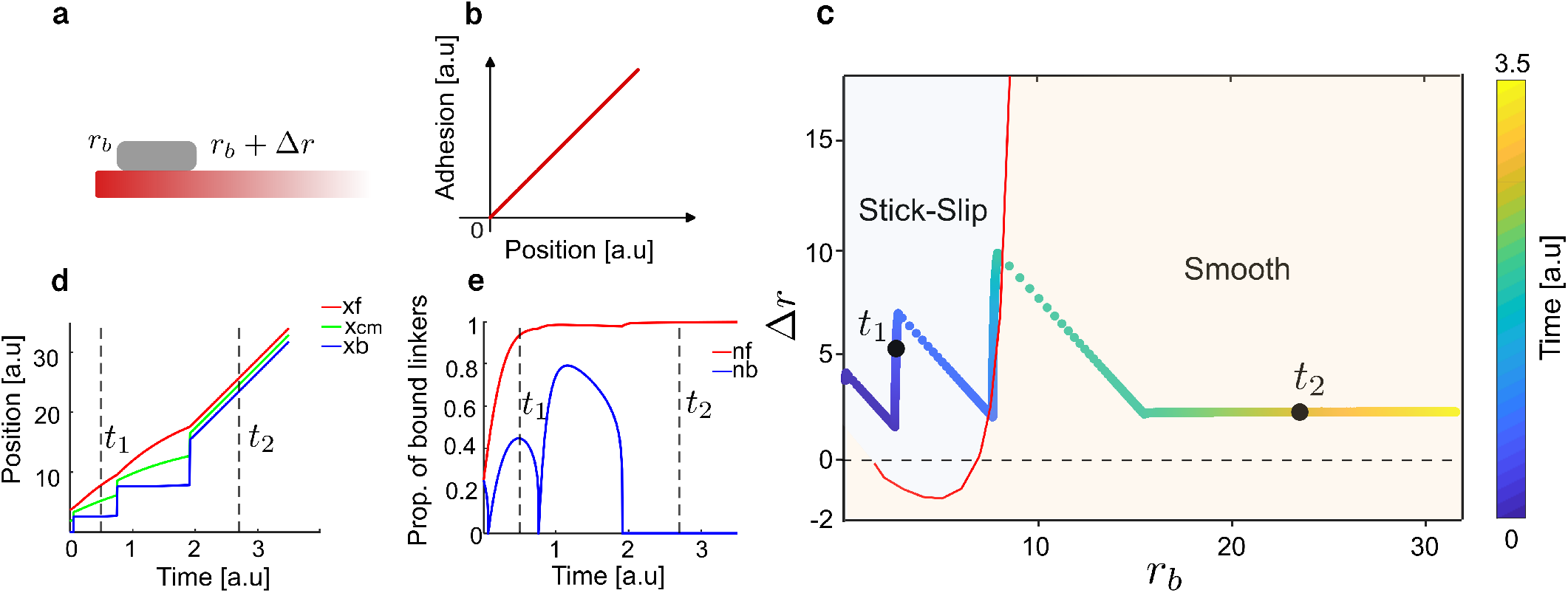
Simulated cell migration on a linear adhesion profile. **a**) Schematic of a cell migrating up the gradient track. **b**) Linearly increasing adhesion profile. **c**) Δ*r* vs *r*_*b*_ phase space diagram for a cell going up the linear gradient. The red line denotes the bifurcation transition between smooth and stick-slip migration, and the colored line denotes the trajectory of the cell shown in the kymograph of (**d**). **e**) The time series of proportion of bound linkers. Dashed lines mark times (*t*_1_, *t*_2_) = (0.8, 3.7). Adhesion parameters: linear profile with *A* = 15 and *k*_*g*_ = 1. Cell parameters: *β*_0_ = 6, 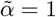 , and *σ*_*v*_ = 0.001.

An example of a cell trajectory kymograph is shown in Fig. 3d, with the corresponding trajectory on the phase diagram in Fig. 3c. As the cell moves up the gradient, increasing *r*_*b*_ drives the trajectory across this boundary, producing a transition from intermittent stick–slip motion to continuous smooth migration. The corresponding internal actin flows at the cell edges and the density of adhesion bound linkers are shown in Fig. 3(e,f). This analysis of the migration on the linear gradient provides a simple dynamical reference for the more complex adhesion profiles considered below.

### 2. Exponential adhesion gradient

Next, we consider migration on a double-exponential adhesion profile (Fig.1b, Eq. (11)) which is the shape was used in the experiments [14]. By varying the activity parameter *β*_0_ we are able to classify four migration modes of cells on this adhesion pattern (Fig. 4). At low activity, *β*_0_ = 2 (Fig. 4A), the cell slowly climbs the gradient but becomes trapped at the adhesion peak, where its position, actin flow, and bound-adhesion linker dynamics relax to steady values. At slightly higher activity parameter, *β*_0_ = 2.5 (Fig. 4B), the cell crosses the peak and enters a smooth oscillatory mode, repeatedly reversing direction around *x*_peak_ = 0 with smoothly alternating actin-flow and adhesion signals. At moderate activity, *β*_0_ = 6.5 (Fig. 4C), the cell displays asymmetric dynamics across the peak: it migrates up the gradient through stick–slip motion, but after crossing the peak it transitions to smooth migration while moving down the gradient. This transition is visible in the position trajectories and is accompanied by a change in actin-flow and bound-linker dynamics from oscillatory to nearly steady behavior. At higher activity, *β*_0_ = 9 (Fig. 4D), there is stick–slip motion on both the ascent and descent of the adhesion gradient. These four migration modes remain robust even at high noise levels (Supplementary Fig. S1).

**Figure 4.**
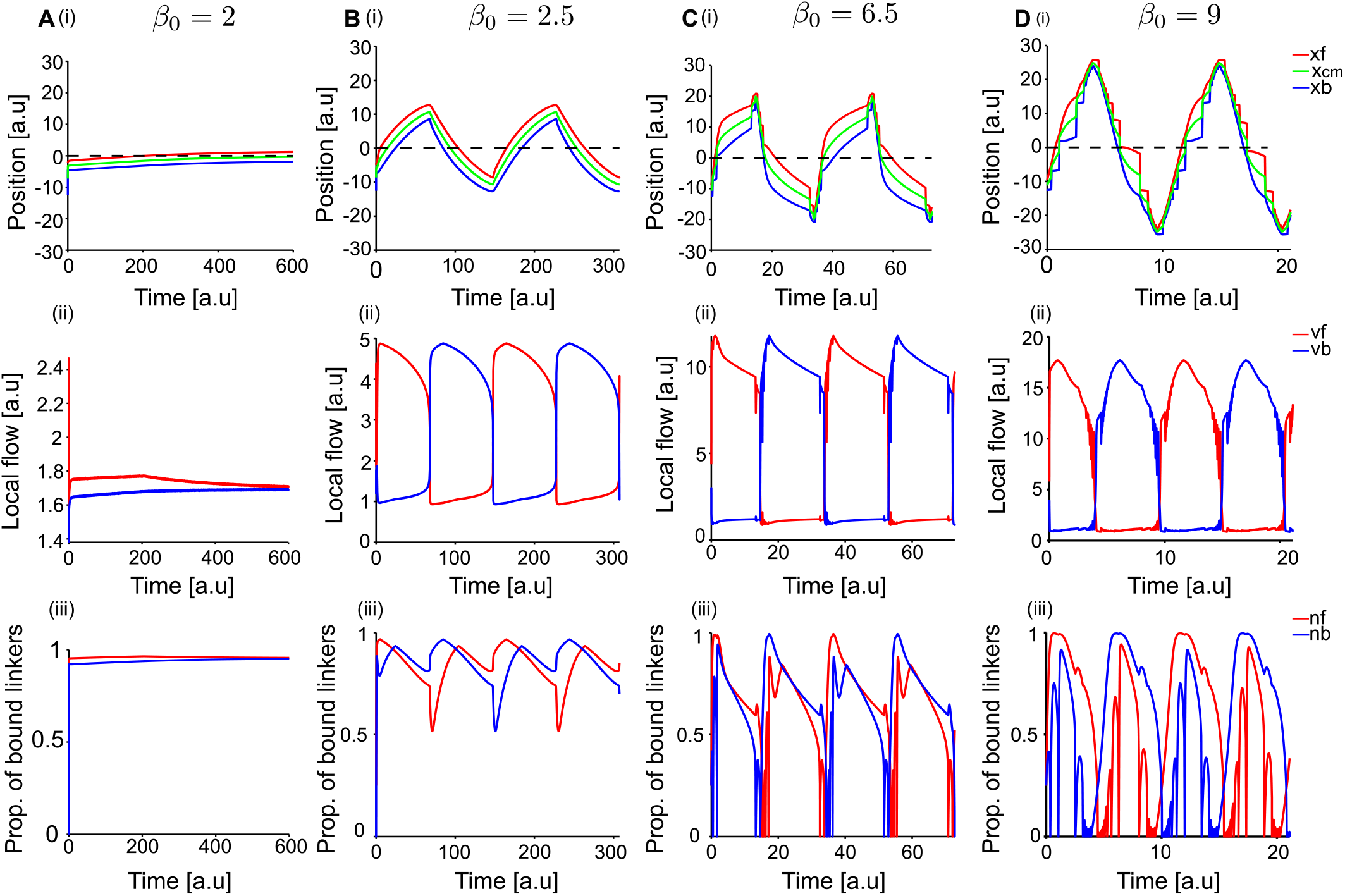
Four modes of migration on an exponential adhesion gradient. **a**[(i)-(iii)] Time series plots at *β*_0_ = 2 showing the cell being trapped at the peak. (i) Position time series: front (red), center of mass (green), and back (blue) of the cell; dotted line indicates the peak of the adhesion gradient. (ii) Local actin flow at the front (red) and back (blue). (iii) Proportion of bound linkers at the front (red) and back (blue). **B**[(i)-(iii)] Time series plots at *β*_0_ = 2.5 showing smooth oscillatory migration around the peak. **C**[(i)-(iii)] Time series plots at *β*_0_ = 6.5 showing stick-slip migration while going up the gradient and smooth migration while moving down the gradient. **D**[(i)-(iii)] Time series plots at *β*_0_ = 9 showing stick-slip migration on both sides of the adhesion peak. Cell parameters: *σ*_*v*_ = 0.01; the panel-specific values of *β*_0_ are indicated above.

The oscillatory migration of the cell around the adhesion maximum predicted by the model (Fig. 4B-D) can be explained as follows. It is not immediately obvious why a cell would overshoot the adhesion maximum and reverse rather than simply arrest there. The answer lies in the adhesion-dependent amplification itself: as the cell approaches the peak, the rising local adhesion progressively boosts protrusive activity, carrying the cell past the maximum. On the far side, adhesion drops, actin protrusive activity falls below the level needed to sustain outward migration, and the cell reverses. The direction-reversal mechanism is therefore activity-based — a direct consequence of the positive adhesion–actin polymerization coupling. At the higher activity regime (Fig. 4C,D) the direction change is associated with a stick-slip event, which can by itself trigger spontaneous directional reversal [16, 33].

To interpret the mixed stick–slip/smooth dynamics observed in Fig. 4C, we examine the dynamics of the cell along the (*r*_*b*_, Δ*r*) phase diagram for *β* = 6 (Fig.5). The phase-space trajectory, shown for very low noise *σ*_*v*_ = 0.001, captures the transition between the different migration regimes on the ascent and descent of the adhesion gradient. The corresponding behavior at higher noise is shown in Supplementary Fig. S2.

**Figure 5.**
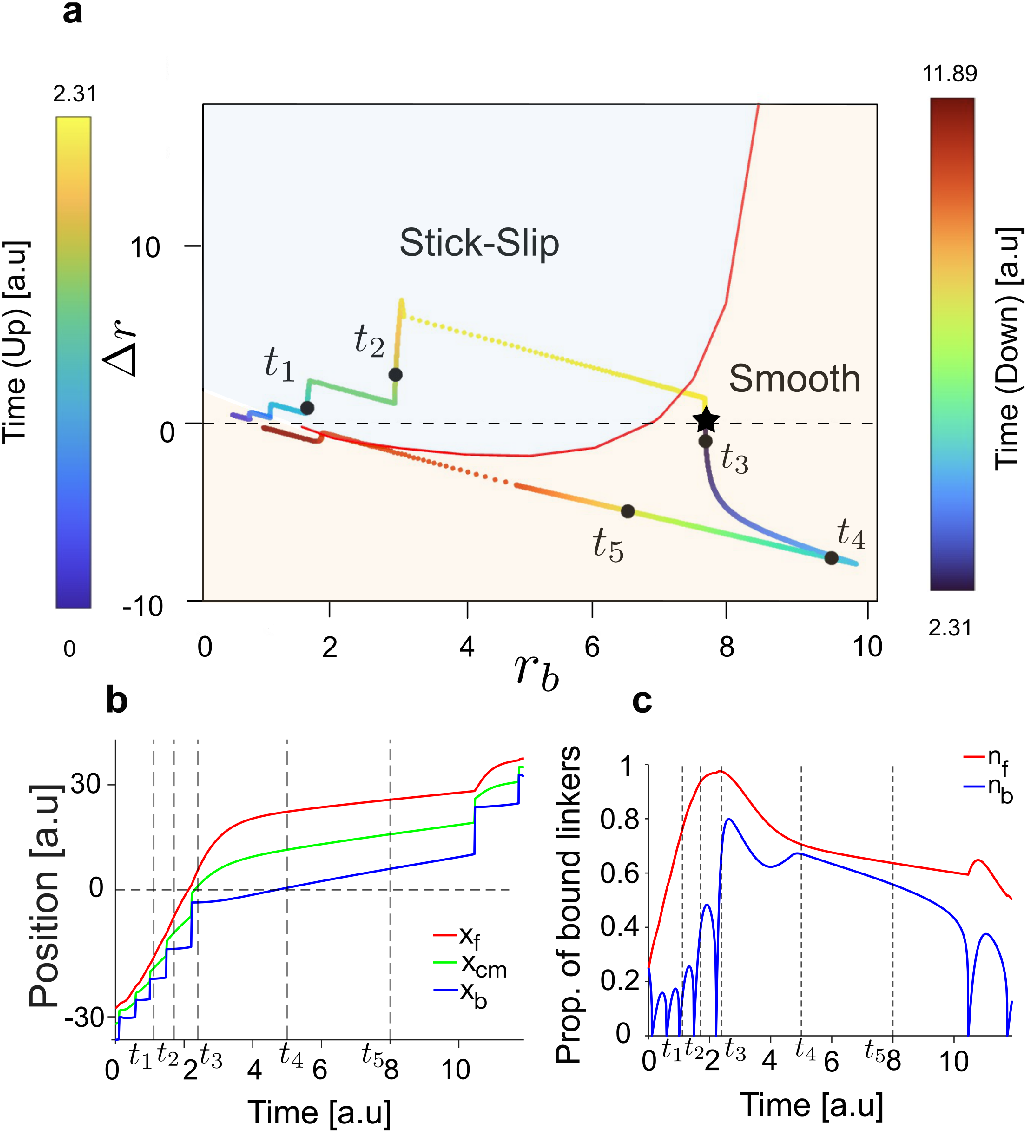
Phase diagram at *β*_0_ = 6. **a**) Δ*r* versus *r*_*b*_ phase diagram, where a time series trajectory is shown (corresponding kymograph in (b)) as a colour gradient from blue (*t* = 0) to yellow (*t* = 2.31) as the cell moves up the gradient, and from blue (*t* = 2.31) to red (*t* = 11.89) as the cell moves down the gradient. Stick-slip and smooth migration regimes are marked, and the transition line is in red. Times *t*_1_ = 1.11, *t*_2_ = 1.7, *t*_3_ = 2.4, *t*_4_ = 5, and *t*_5_ = 8 are marked in both the phase space diagram and subsequent time series plots, showing stick-slip mode at *t*_1_ and *t*_2_, and smooth mode at *t*_3_, *t*_4_, and *t*_5_. The point at which the cell crosses the peak is marked with a black star. **b**) Position time series plot with marked times shown as vertical dotted lines. The position at which the adhesion value becomes maximum is marked by a horizontal dotted line at *y* = 0. Positions of the front, center, and back of the cell are shown in red, green, and blue, respectively. **c**) Proportion of bound linkers time series plot with marked time points shown as vertical dashed lines. Front and back values are shown in red and blue, respectively. Cell parameters: *β*_0_ = 6 and *σ*_*v*_ = 0.001.

### B. Experimental Comparison

#### 1. Validation of haptotaxis model based on adhesion-driven cell polarization

To validate the one-dimensional model, we compared its predictions with experiments on fibronectin-coated one-dimensional tracks containing a prescribed adhesion gradient [14]. Representative kymographs from experiments and simulations show comparable migration patterns relative to the peak-adhesion position at *x*_peak_ = 0, and individual trajectories display similar cell-to-cell variability in both datasets (Fig. 6**a–d**).

**Figure 6.**
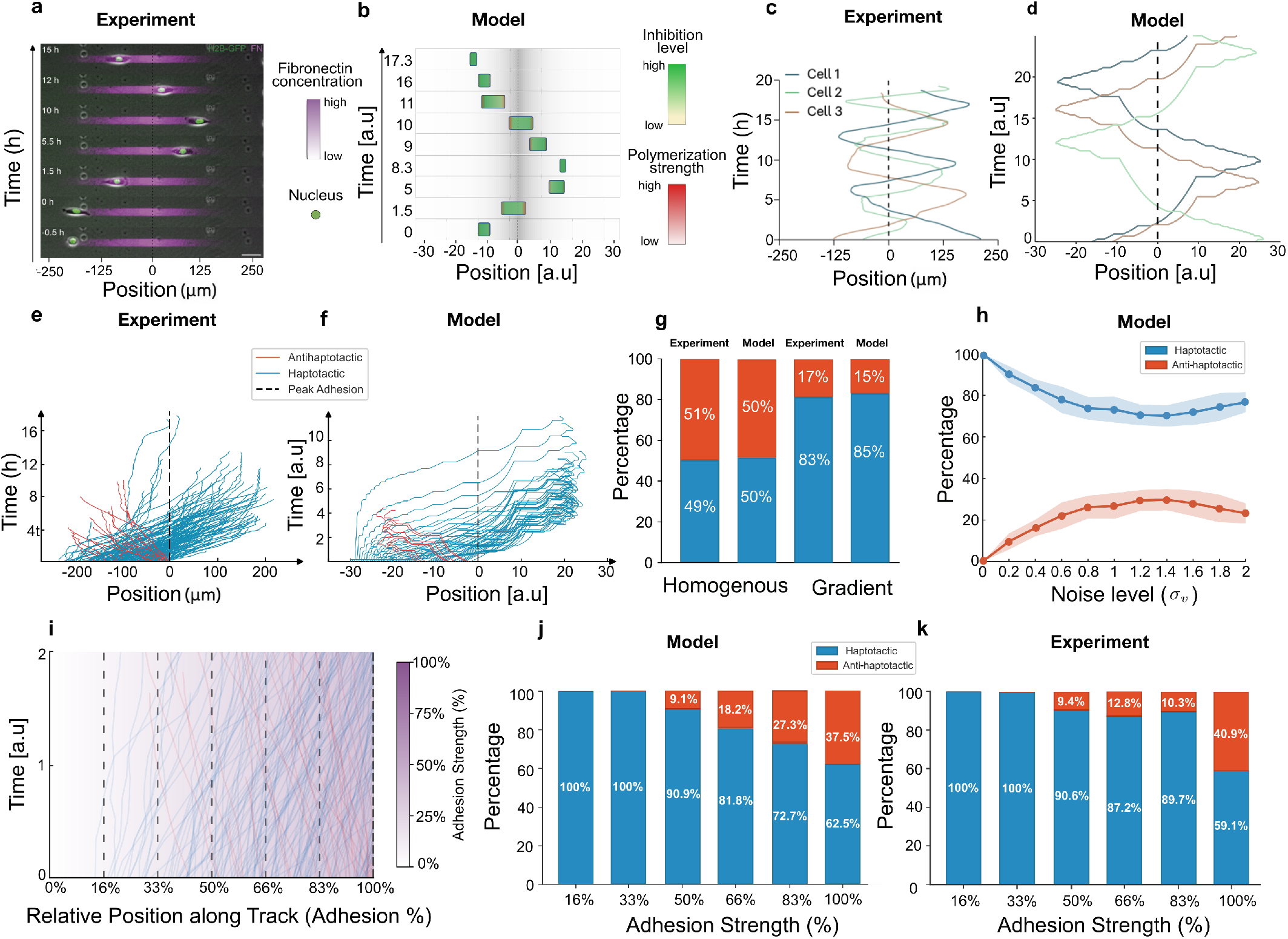
Quantitative comparison between experiments and simulations on 1D adhesion-gradient tracks. **a**) Representative experimental snapshots showing the cell position as a function of time on a 1D track, coated with a spatial gradient of fibronectin concentration (magenta intensity). **b**) Corresponding simulation snapshots using the 1D model under the same track geometry, showing the cell position over time. **c**) Representative experimental trajectories of three individual cells migrating on adhesion-gradient tracks. **d**) Representative simulated trajectories (three realizations) obtained from the 1D model. The dashed vertical line, at *x* = 0, denotes the peak-adhesion position. **e**) First-run experimental trajectories on adhesion gradients (*N* = 173), classified by directionality: *haptotactic* (blue; motion toward higher adhesion) and *anti-haptotactic* (orange; motion toward lower adhesion). **f**) First-run simulated trajectories displayed and classified as in (**e**). **g**) Proportions of haptotactic trajectories (blue) for homogeneous versus gradient adhesion conditions, comparing experiments and simulations. Experimental sample sizes: homogeneous (*N* = 202) and gradient (*N* = 173) fibronectin patterns. **h**) Simulation results showing the dependence of directionality on noise amplitude *σ*_*v*_ : the haptotactic fraction decreases with increasing noise, while the anti-haptotactic fraction increases (shaded regions denote variability across realizations). **i**) Position resolved haptotactic migration. Positions are binned uniformly along the track and plotted as a percentage of the distance from the gradient start (0%) to the peak-adhesion location (100%). **j**) Position-resolved directionality from the model, computed using the same binning procedure as in (**i**). **k**) Corresponding position-resolved directionality in experiments: percentages of haptotactic (blue) and anti-haptotactic (red) trajectories as a function of initial position along the gradient. Cell parameters: *β*_0_ = 9 in panel **d**, *β*_0_ = 6.5 in panels **j** and **i**. Noise amplitudes are *σ*_*v*_ = 0.1 in panel **d**, *σ*_*v*_ = 0.6 in panel **f** and *σ*_*v*_ = 0.4 in panel **j**. Simulation durations are *T* = 25 in panel **d**, *T* = 18 in panel **f**, and *T* = 12 in panel **j**.

In these comparisons we used model parameters which were previously calibrated to fit the behavior of motile cells of different types [16, 30, 31]. We chose the adhesive pattern length in the model units to be ≈3 − 5 times the average cell length, to correspond to the experimental conditions (Fig. 6**a,b**). Note that all lengths in the simulations are in units of the un-spread length of the cell [16]. Representative trajectories of the cell’s center-of-mass in both experiments and simulations are shown in Fig. 6**c,d**.3

To quantify directional bias, we analyzed the first-run segment of each trajectory—defined as the motion from the initial position until the first reversal or effective stop—classifying runs as haptotactic if the net displacement was toward increasing adhesion, and anti-haptotactic otherwise (Fig. 6**e,f**). With the right set of parameters the model quantitatively reproduces the observed bias: on gradient tracks, 83% of experimental and 85% of simulated trajectories are haptotactic, whereas homogeneous tracks show no preference (49% and 50%, respectively), confirming that directionality arises from the adhesion gradient rather than intrinsic polarity bias (Fig. 6**g**). Varying the velocity-noise amplitude in the simulations (*σ*_*v*_) shows that haptotaxis remains robust (*>* 80%) at low noise but degrades at high noise (Fig. 6**h**), which overwhelms the adhesion bias on the polarity (Eq.1). This dependence also varies with the baseline activity *β*_0_ (SI Fig.S3).

Furthermore, the model predicts that the haptotactic directionality decreases as cells start closer to the adhesion peak (Fig. 6**i,j**), due to the reduction in Δ*β* across the cell near the peak, which weakens front–rear polarization and lowers the directionality (SI Fig.S3). The predicted trend is verified in the experimental data (Fig. 6**k**).

Together, these results demonstrate that adhesion-dependent modulation of protrusive activity at the cell’s leading-edges is sufficient to reproduce the experimentally observed haptotactic bias, and its dependence on the initial position along the gradient.

The model reproduces position-dependent trends in cell morphology and kinematics (Fig. 7). Haptotactic cells elongate as they approach the adhesion maximum and migrate fastest up the gradient, with velocity decreasing progressively during down-gradient motion [14]. In the model, these trends arise from the position dependence of the front–back protrusive asymmetry, which is strongest while approaching the peak and weakens after crossing it, simultaneously reducing mean speed and elongation (Fig. 7**a–d**).

**Figure 7.**
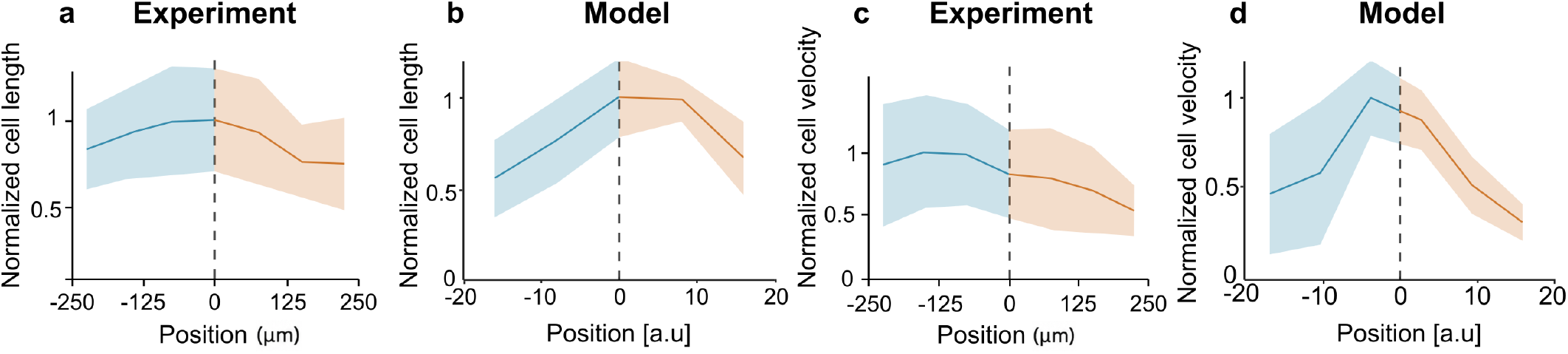
Position-dependent cell length and velocity: experiment–model comparison. Cell length and velocity are plotted as functions of position along the adhesion-gradient track. These quantities are normalized by their maximal value. The peak-adhesion position is indicated by the vertical dashed line at *x*_peak_ = 0. Blue curves correspond to the portion of the first runs (as in Fig.6e,f) migrating *up* the gradient (approaching the peak), and orange curves correspond to the portion *down* the gradient (after crossing the peak). Shaded regions denote respective errors. **a**) Experimental, and **b**) model mean cell length, normalized by the maximal value. **c**) Experimental and **d**) model mean cell speed, normalized by the maximal value. Cell parameters: *β*_0_ = 7.5 and *σ*_*v*_ = 0.4. Statistics are computed from *N* = 2000 different simulated trajectories.

We next tested whether the model reproduces the effects of blebbistatin-induced myosin II inhibition. In our coarse-grained model, cellular contractility governs the mechanical coupling between cell edges, and we therefore represent blebbistatin treatment by reducing the spring stiffness from *k* = 0.8 to *k* = 0.4, which was previously shown to capture qualitatively the effects of myosin-II inhibitory drugs [30]. This single change reproduces all key experimental observations [14]: the haptotactic bias on the adhesion gradient tracks is maintained (Fig. 8**a,b**), cell length distributions shift to larger values (Fig. 8**c,d**), and speed distributions shift toward lower values (Fig. 8**e,f**), consistent with slower, more elongated cells under reduced contractility.

**Figure 8.**
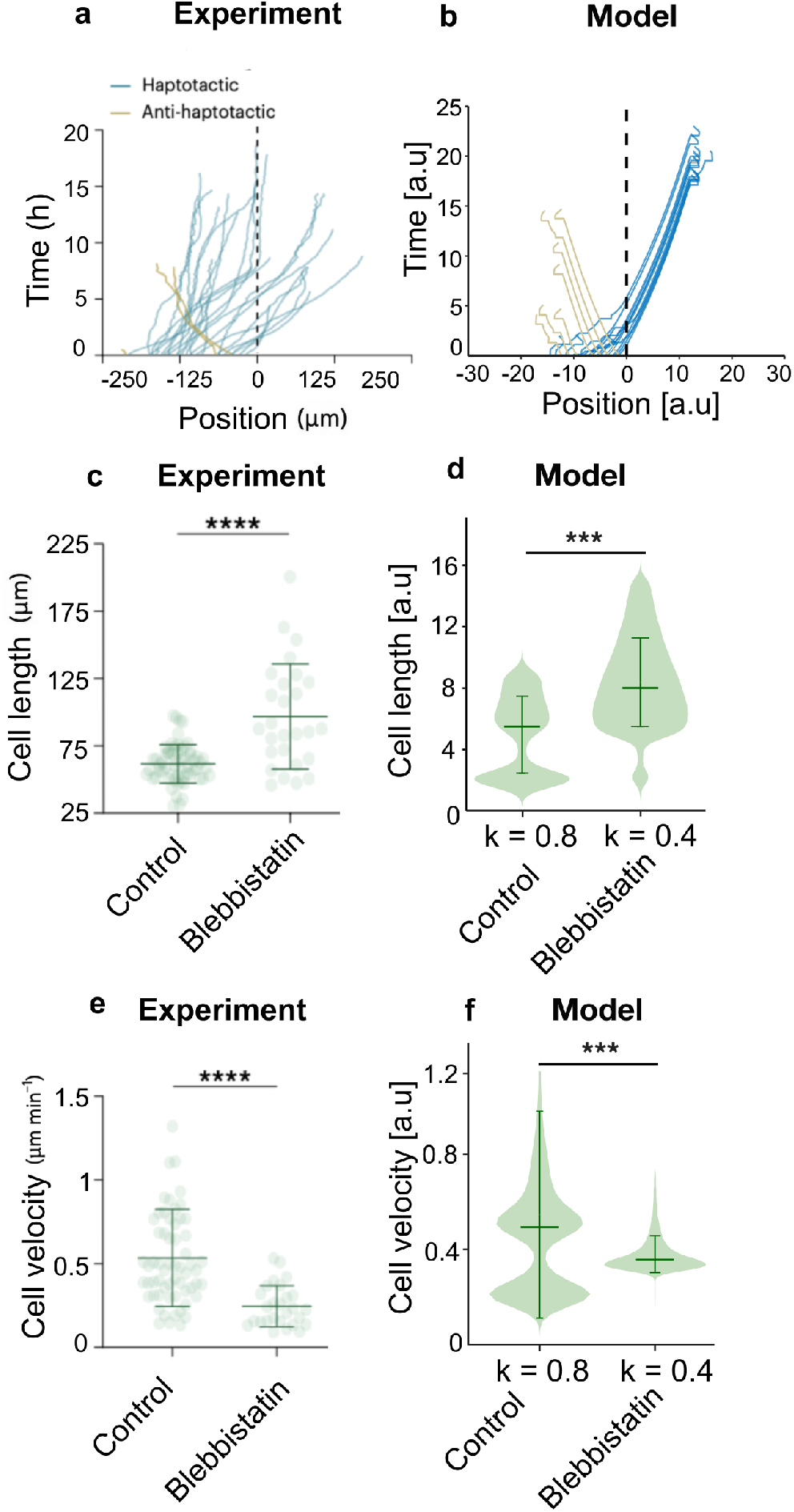
Cell migration on adhesion gradient under blebbistatin treatment. **(a–b)** First-run trajectories from experiments with blebbistatin treatment (N = 27) and corresponding simulations with reduced spring constant (from *k* = 0.8 to *k* = 0.4). **(c–d)** Mean cell length under blebbistatin treatment compared to control (N = 61 control, N = 27 blebbistatin for experiments), and corresponding simulation results with reduced spring constant. **(e–f)** Mean velocity for blebbistatin-treated versus control cells in experiments, and corresponding simulation results with reduced spring constant. For panels (**c**) and (**e**), significant differences between groups were identified using the Kruskal–Wallis test, with pairwise comparisons performed using Dunn’s multiple comparisons test (\*\*\*\**P <* 0.0001). For panels (**d**) and (**f**), significant differences were identified using the Mann– Whitney *U* test (\*\*\**P <* 0.001). Error bars represent standard deviation. Cell parameters: *β*_0_ = 6.5; for panels **b, d**, and **f**, *σ*_*v*_ = 0.4 and *T* = 25.

The adhesion-gradient magnitude was shown in experiments to affect the strength of the directional bias [14], which we therefore tested in our model. The experimental haptotactic fraction increases systematically from 73% to 83% to 98% as the concentration of surface ECM was increased (Fig. 9 **a**). Similarly, the model reproduces this trend (Fig. 9**b**) for increasing values of the adhesion magnitude (*A* in Eq.11). The corresponding speed statistics shows the same qualitative agreement between experiment and model (Fig. 9 **c,d**). Additionally, the model predicts systematic changes in cell length with increasing adhesion amplitude, reported in Supplementary Fig. S4, providing an independent morphological read-out consistent with the gradient-magnitude dependence of the migration response.

**Figure 9.**
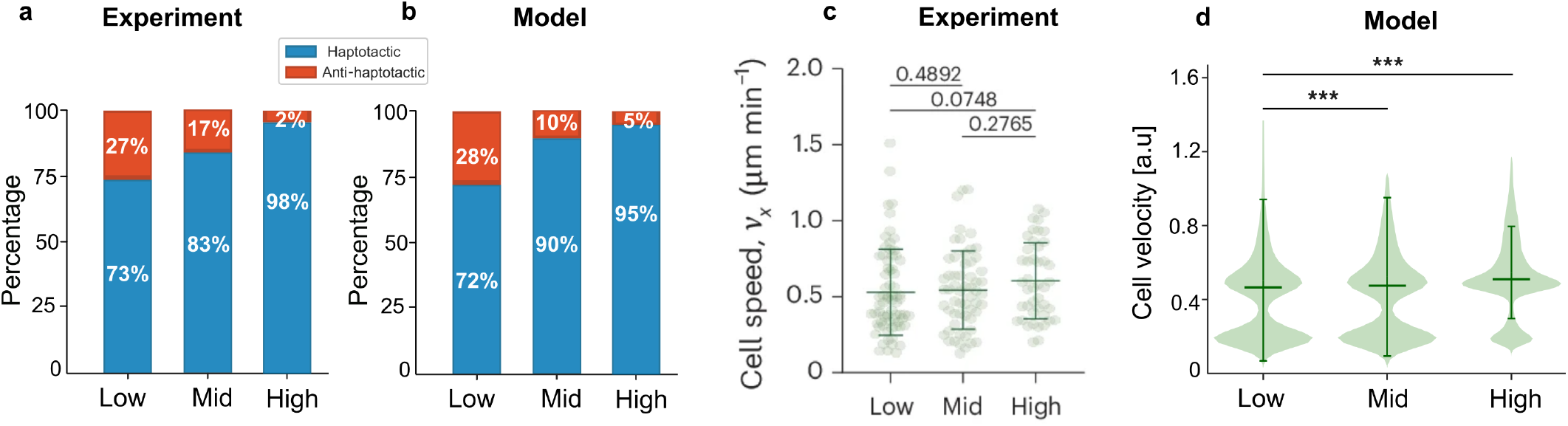
Effect of adhesion-gradient magnitude: experiment–model comparison. **a**) Experimental percentages of haptotactic (blue) and anti-haptotactic (red) first-run trajectories for low (N = 71), mid (N = 58), and high (N = 44) 1D fibronectin concentration gradient magnitudes. Low, mid and high magnitudes have mean fibronectin densities of 24.293±10.244 ng*/*cm^2^, 46.784±12.82 ng*/*cm^2^ and 67.579±20.613 ng*/*cm^2^, respectively. **b**) Corresponding simulation percentages for low, mid, and high adhesion gradient magnitudes. **c**) Experimental peed distributions of cells migrating on low (N = 71), mid (N = 58) and high (N = 44) 1D fibronectin gradient magnitudes. Error bars are s.d., P values were obtained using the Mann– Whitney test. **d**) Model speed distributions for the same conditions. Error bars represent standard deviation, significant differences were identified using the Mann–Whitney *U* test (\*\*\**P <* 0.001). Cell parameters: *β*_0_ = 7.5 and *σ*_*v*_ = 0.4. Low, mid and high adhesion gradients in the simulations correspond to maximal values of the parameter *A* = 22, 27, 32 respectively (Eq.11). Statistics are computed from *N* = 2000 simulated trajectories.

Finally, we examined cell migration on an inverted adhesion profile, where adhesion is minimal at the track center (*x* = 0) and increases toward the ends (Fig. 10 **a,b**). First-run trajectories (Fig. 10 **c,d**) were classified as haptotactic or anti-haptotactic relative to the local adhesion gradient. In both experiment [14] and model, the majority of first runs were directed toward increasing adhesion (Fig. 10 **e**), demonstrating that directional bias is governed by local gradient sensing rather than an absolute preference for the track center. These results confirm that the model can explain gradient-dependent directionality across general substrate geometries.

**Figure 10.**
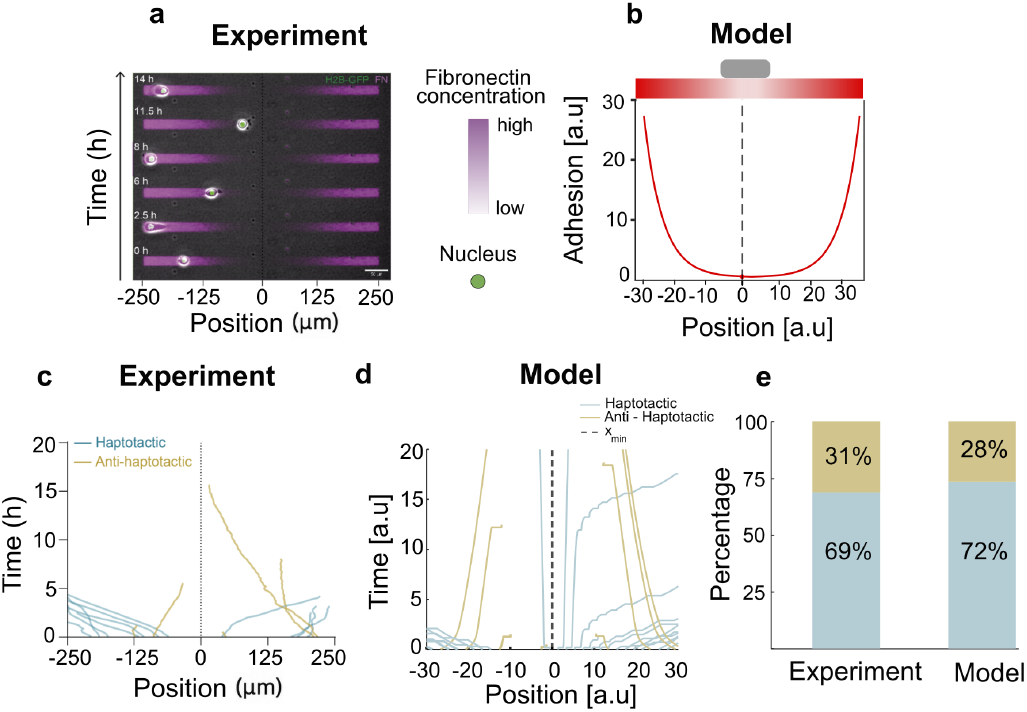
Migration on an inverted adhesion profile: experiment–model comparison. **a**) Experimental snap-shots on an inverted adhesion profile (adhesion is minimal at the center and increasing toward the ends). **b**) Corresponding simulation schematic showing the inverted profile. **c**,**d**) Experimental and simulated first-run trajectories classified as haptotactic or anti-haptotactic. **e**) Percentages of haptotactic and anti-haptotactic first runs in the experiment and model (corresponding to the first-runs shown in (c,d)). Cell parameters: *β*_0_ = 6 and *σ*_*v*_ = 0.4.

### 2. Variability of cell migration patterns and intrinsic activity

The comparisons we showed so far between the model and the experiments used a narrow range of parameters. In particular, we used a value of *β*_0_ ∼ 7 which determines the actin polymerization activity and controls the cell speed. However, experimental data indicates that there is a large variability between cellular migration speeds. Haptotactic and anti-haptotactic trajectories are shown separately in Fig. 11 **a,c**, with their corresponding speed distributions in Fig. 11 **b,d**. This suggests that the value of *β*_0_ has a relatively wide distribution in the cell population.

**Figure 11.**
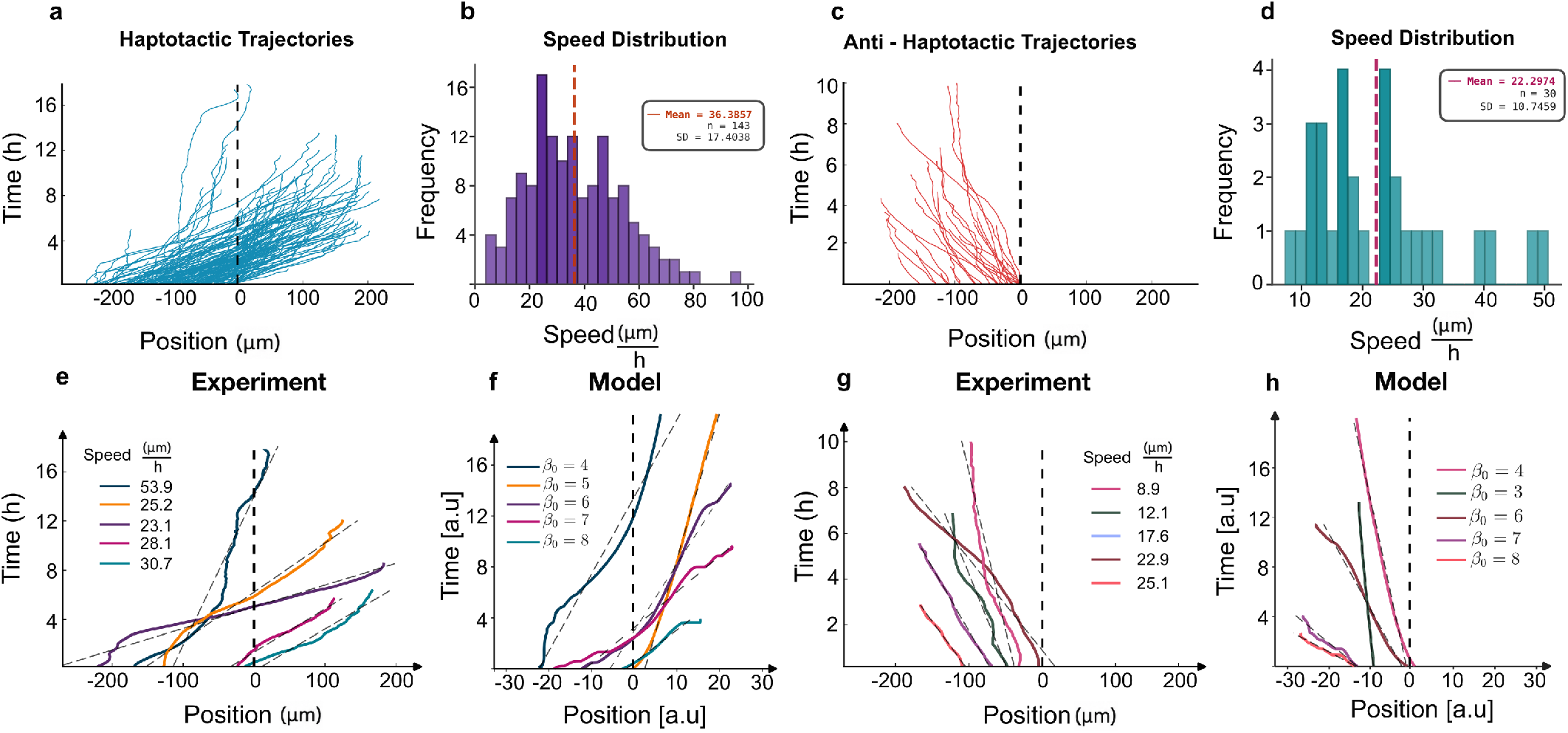
Experimental speed distributions and model correspondence. **a**) Experimental haptotactic first-run trajectories, and (**b**) their corresponding speed distribution. **c**) Experimental anti-haptotactic first-run trajectories, and (**d**) their corresponding speed distribution. **e**) Examples of experimental haptotactic first-run trajectories, with dashed lines denoting their mean migration speeds (values indicated in legend). **f**) Simulated haptotactic trajectories at different migration speeds obtained by varying *β*_0_ (legend). **g**) Experimental anti-haptotactic first-run trajectories, with dashed lines denoting their mean migration speeds (values indicated in legend). **h**) Simulated anti-haptotactic trajectories at different migration speeds obtained by varying *β*_0_ (legend). Cell parameters: *σ*_*v*_ = 0.4; different simulated migration speeds are obtained by varying *β*_0_, as indicated in the legend.

To compare this variability with the model, representative trajectories with similar slopes were grouped by color and compared with simulations obtained for different values of the activity parameter *β*_0_ Fig. 11 **e–h**. The qualitative agreement suggests that variations in activity, captured by ∼2 *< β*_0_ *<*∼9, can account for the experimentally observed spread in migration speeds in both haptotactic and anti-haptotactic cells. Our model predicts that different values of the cellular activity correspond to different migration patterns on the adhesion gradients (Fig.4). These predicted migration modes are indeed observed in experiments (Fig. 12): trapping near the peak-adhesion region (**a**), smooth continuous migration (**b**), mixed dynamics with stick–slip up-gradient and smooth motion down-gradient (**c**), and predominantly stick–slip migration throughout (**d**). Across all four modes, the model reproduces the qualitative kymograph features observed experimentally, providing a trajectory-level validation that complements the population-averaged comparisons shown above. Note that in addition to the different values of the activity parameter *β*_0_, the noise also parameter *σ*_*v*_ also plays an important role (for example see Fig. 12a,b).

**Figure 12.**
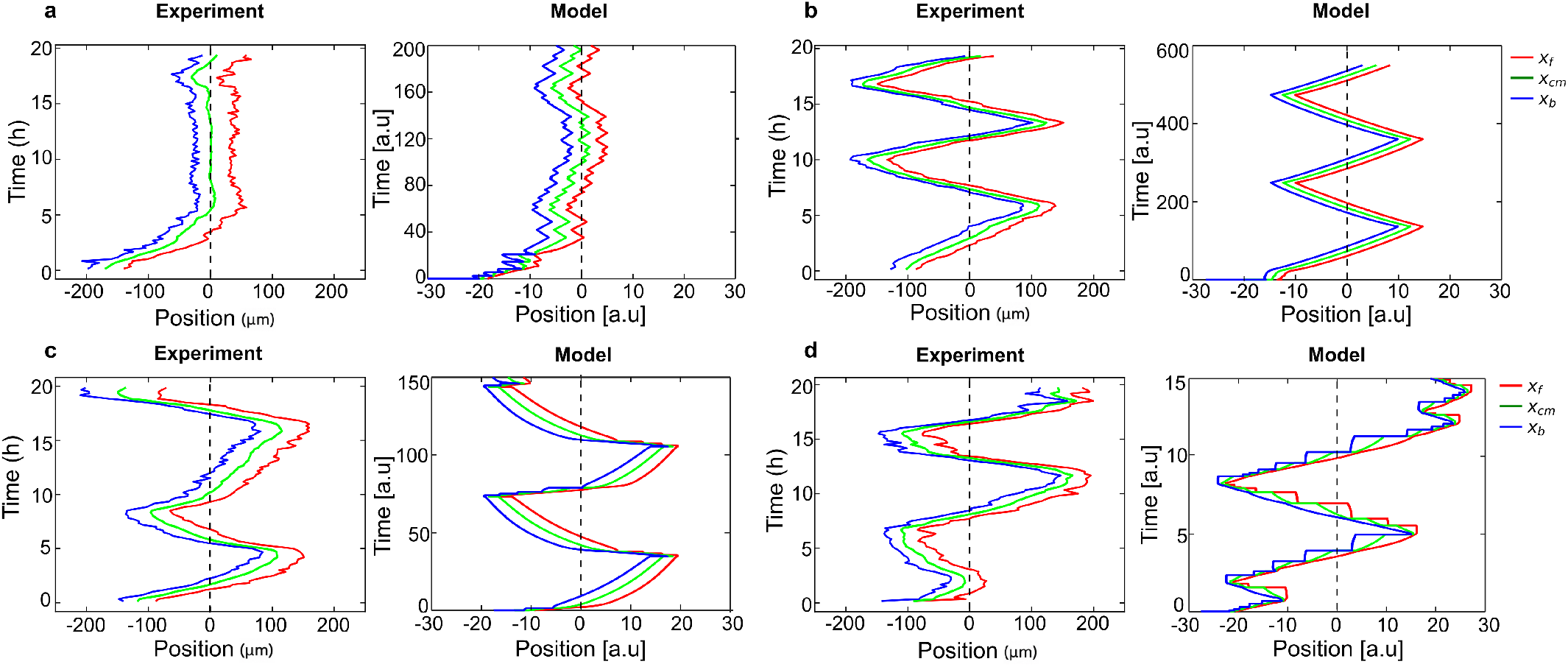
Kymograph comparisons of representative single-cell trajectories. Each panel shows an experimental kymograph (left) and a corresponding model trajectory (right). The vertical dashed line indicates the peak-adhesion position (*x*_peak_ = 0). Red, green, and blue curves denote the right edge *x*_*f*_ , cell center *x*_cm_, and left edge *x*_*b*_, respectively. **a**) A cell getting trapped near the peak: the trajectory remains localized in the vicinity of the peak-adhesion region. **b**) Smooth migration: continuous motion without pronounced intermittent arrest. **c**) Mixed dynamics: stick–slip during the migration up the gradient, and smoother motion during the migration down the gradient. **d**) Predominantly stick–slip migration. Cell parameters: **a**, *T* = 200, *β*_0_ = 2.4, *σ*_*v*_ = 0.8, *r*_0_ = 0.15, 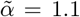, *x*_*f*_ = 12, *x*_*b*_ = 2; **b**, *T* = 500, *β*_0_ = 1.95, *σ*_*v*_ = 0.2, *r*_0_ = 0.15, 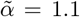, *x*_*f*_ = 12, *x*_*b*_ = 5; **c**, *T* = 150, *β*_0_ = 5.0, *σ*_*v*_ = 0.2, *r*_0_ = 0.75, 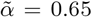, *x*_*f*_ = 23, *x*_*b*_ = 15; **d**, *T* = 15, *β*_0_ = 9.0, *σ*_*v*_ = 0.675, *r*_0_ = 0.55, 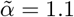, *x*_*f*_ = 11, *x*_*b*_ = 5.

Next we compare different cellular trajectories Fig. 13, where different migration modes are reproduced in the simulations by varying the cell activity *β*_0_, and the actin flow noise *σ*_*v*_ parameters. In Fig. 14 we plot these examples in the corresponding (*β*_0_, *σ*_*v*_) parameter space diagram.

**Figure 13.**
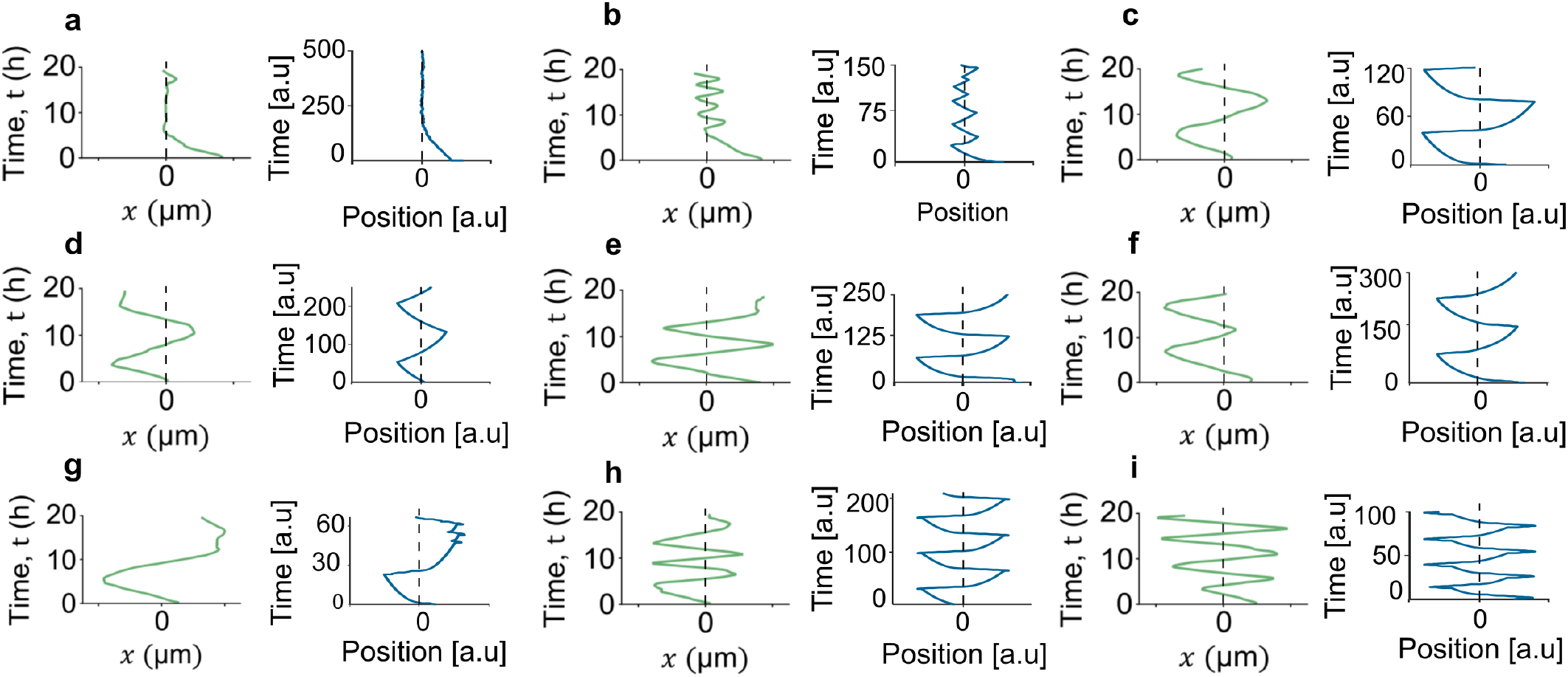
Representative oscillatory trajectories: experiment–model comparison. For each panel, an experimental trajectory (green) and a representative model trajectory (blue) are shown as position versus time, illustrating oscillatory motion around the peak-adhesion region (vertical dashed line *x* = 0). The simulations are done by varying only the cell activity *β*_0_, and the actin flow noise *σ*_*v*_ parameters. Cell parameters: **a**, *β*_0_ = 2, *σ*_*v*_ = 0.2; **b**, *β*_0_ = 2.2, *σ*_*v*_ = 1; **c**, *β*_0_ = 2.5, *σ*_*v*_ = 0.5; **d**, *β*_0_ = 2.5, *σ*_*v*_ = 0.2; **e**, *β*_0_ = 3.5, *σ*_*v*_ = 0.4; **f**, *β*_0_ = 3.5, *σ*_*v*_ = 0.2; **g**, *β*_0_ = 5, *σ*_*v*_ = 1; **h**, *β*_0_ = 5, *σ*_*v*_ = 0.2; **i**, *β*_0_ = 5.5, *σ*_*v*_ = 0.1.

**Figure 14.**
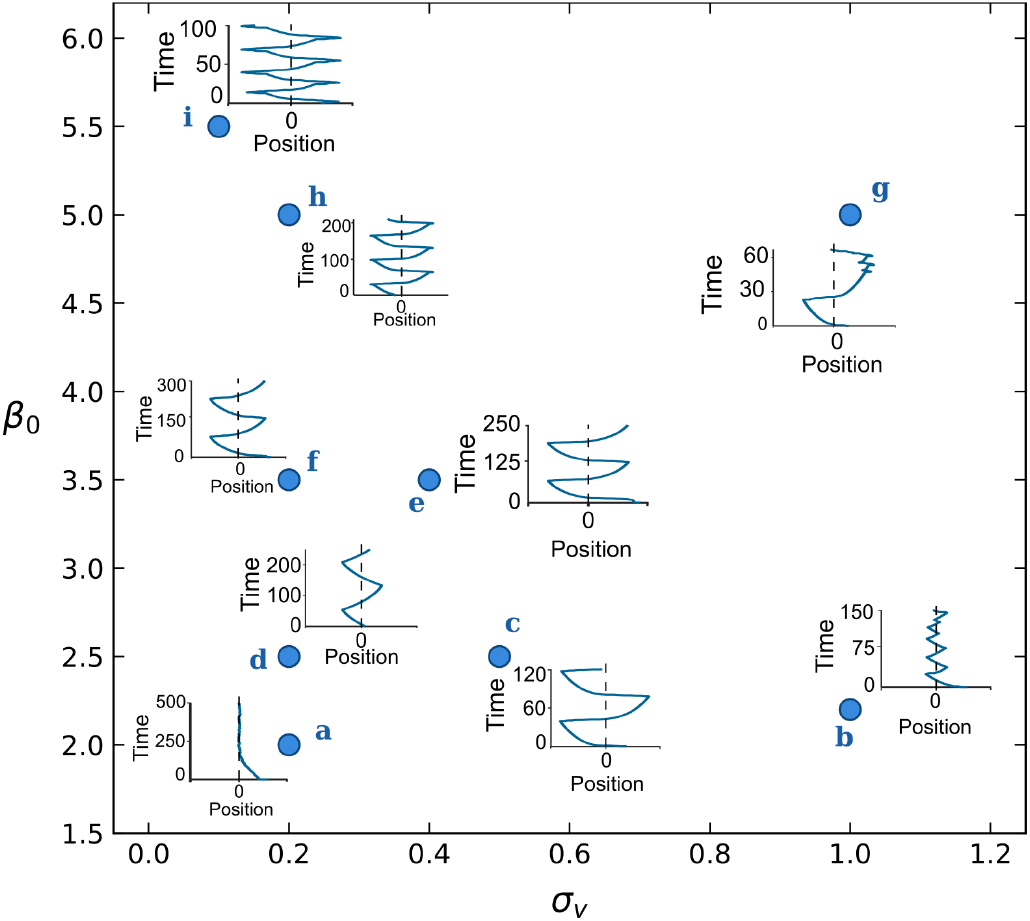
Parameter phase space diagram for the simulated trajectories shown in Fig.13, as function of the cell activity (*β*_0_) and noise (*σ*_*v*_) parameters, that span the range of observed trajectories.

Finally, we compared experimental and model traction kymographs across the four migration modes (Fig. 15). The comparison is intentionally *qualitative*, assessing whether the model reproduces the characteristic spatiotemporal traction forces of each mode rather than matching absolute magnitudes. For further description of the traction force calculation and plotting, see Supplementary Information section S5.

**Figure 15.**
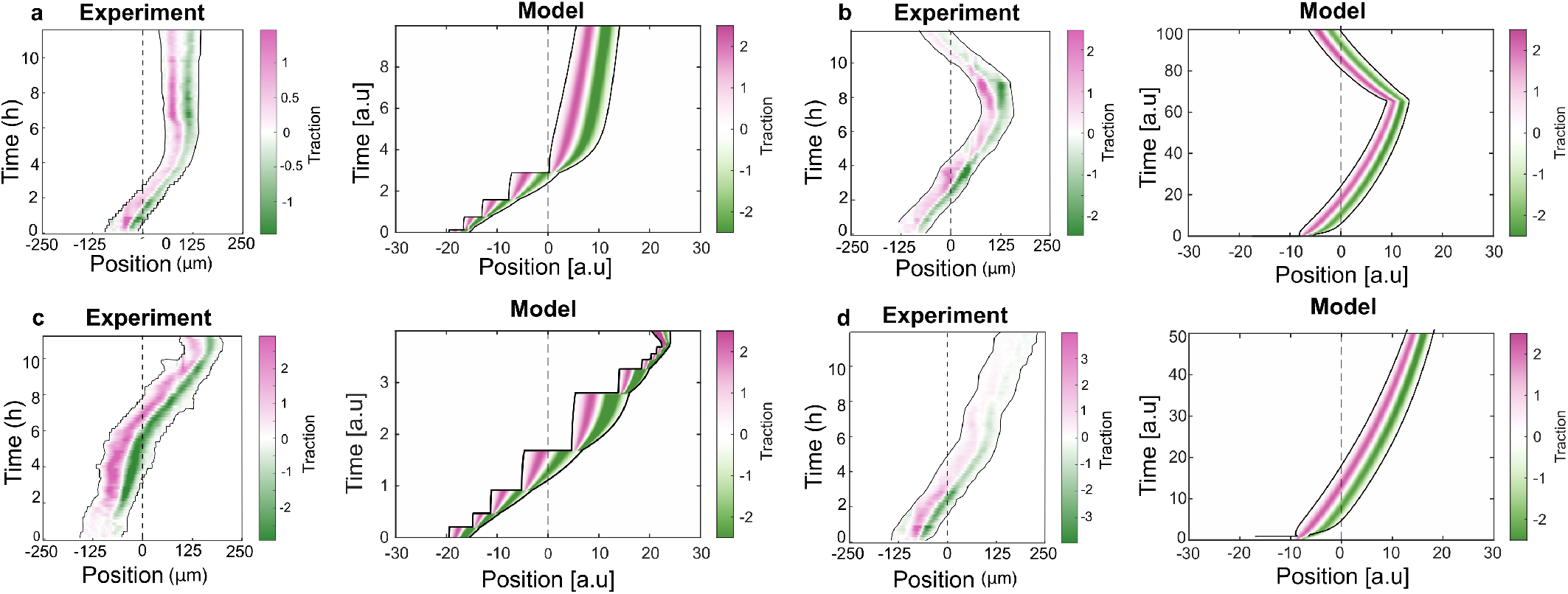
Traction kymograph comparison between experiments and model. For each panel, the experimental traction kymograph (left) is shown alongside the corresponding model traction field (right), plotted over the same space–time axes. The vertical dashed line indicates the peak-adhesion position (*x*_peak_ = 0). Color encodes the signed traction field (see color bars). Cell parameters: **a**, *T* = 10, *β*_0_ = 5.5, *σ*_*v*_ = 0.3, *r*_0_ = 0.75, 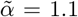, *x*_*f*_ = 23, *x*_*b*_ = 15; **b**, *T* = 100, *β*_0_ = 2.25, *σ*_*v*_ = 0.2, *r*_0_ = 0.35, 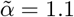, *x*_*f*_ = 12, *x*_*b*_ = 5; **c**, *T* = 3.25, *β*_0_ = 9.0, *σ*_*v*_ = 0.2, *r*_0_ = 0.55, 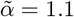, *x*_*f*_ = 11, *x*_*b*_ = 5; **d**, *T* = 50, *β*_0_ = 2.3, *σ*_*v*_ = 0.2, *r*_0_ = 0.15, 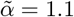, *x*_*f*_ = 11, *x*_*b*_ = 5.

Overall, we demonstrate that the model can capture the whole spectrum of the observed cellular migration patterns on the adhesion gradients. The main origin of this variability is shown to be the variable intrinsic cellular activity (*β*_0_), and the noise *σ*_*v*_, which also controls the overall speed of the cell. On the symmetric adhesion pattern this activity variations give rise to the observed spectrum of migration patterns.

## IV. CONCLUSIONS

We developed a theoretical model for cell migration on confined adhesion gradients by extending a validated one-dimensional framework to substrates with spatially varying adhesion. The essential new ingredient is a local coupling between adhesion strength and actin polymerization and protrusive activity at the leading-edges of the cell. This coupling provides a minimal and biologically motivated mechanism by which spatial differences in adhesiveness across the cell body bias the spontaneous self-polarization mechanism of the cell, that is based on a front–rear asymmetry in protrusive activity. This process biases migration toward higher adhesion without invoking explicit gradient-comparison modules.

When compared quantitatively with experiments, the model captures the enrichment of haptotactic first runs relative to homogeneous controls. In addition, the model predicts that the degree of haptotactic directionality depends on the position along the gradient, which is verified in experiments. It accounts for the the position-dependent trends in cell length and velocity during up- and down-gradient motion. The model reproduces the effects of adhesion strength on the directional bias, as well as the effects of myosin-II inhibition using blebbistatin treatment. Finally, the model reproduces the haptotactic bias under an inverted adhesion profile without any parameter adjustment, confirming that it can describe migration on general adhesion patterns.

At the single-cell level, the model predicts a rich set of migration modes as function of the intrinsic level of cellular actin polymerization and protrusive activity at the leading edge. Since the experimental data exhibits a wide range of cellular speeds, we are able to relate the observed variability in cellular migration patterns to the inherent variability in cytoskeletal activity. The experimentally observed migration patterns on the adhesion gradients match well the model’s predicted spectrum of migration characteristics. Most importantly, by including an explicit model for the spontaneous self-polarization of the cell and coupling it to the adhesion, the model is able to explain why weakly active cells (small *β*_0_) have low persistence and get trapped at the peak of the adhesion pattern (Fig.14). Cells with higher activity (large *β*_0_) are persistent across the adhesion peak, and exhibit oscillatory migration up and down the adhesion gradient (Fig.14).

Taken together, these results establish adhesion-dependent amplification of edge activity, combined with mechanical front–rear coupling and stochastic fluctuations, as a minimal and sufficient physical basis for confined haptotaxis and oscillatory migration on patterned adhesion landscapes. The model offers a quantitative foundation for future work varying gradient shape, amplitude, and confinement geometry, and could be extended to incorporate lateral shape dynamics, traction-dependent adhesion maturation, and the transition to two-dimensional migration. In addition, the model could be used to explore collective haptotaxis of cellular clusters [29].

## V. ACKNOWLEDGEMENT

This work was supported by Lee and William Abramowitz Professorial Chair of Biophysics (N.S.G.). This research is made possible in part by the historic generosity of the Harold Perlman Family (N.S.G.). N.S.G. acknowledges support from the Human Frontier Science Program grant RGP0032/2022 and ISF-DFG grant 1182/25. UR acknowledges the support of DST-INSPIRE Faculty Fellowship (DST/INSPIRE/04/2022/003052), Government of India.X.T. is supported by the European Research Council (ERC Advanced Grant 883739), the Human Frontier Science Program (HFSP RGP022/2024), the Spanish Ministry of Science, Innovation and Universities (PID2021-128635NB-I00), the Generalitat de Catalunya (AGAUR SGR-2021-01425 and CERCA Programme), and the La Caixa Foundation (LCF/PR/HR24/00326). This work was supported in part by the Spanish State Research Agency (AEI; grant PID2024-159132OB-I00 to R.S.). R.S. is a Serra Húnter Fellow.

