## Supplementary Information for "A minimal physical model of adhesion-dependent protrusion explains confined haptotaxis and oscillatory cell migration"

### S1. FOUR MODES OF MIGRATION AT HIGHER NOISE LEVEL ($\sigma_v = 0.4$)

In Fig.S1 we plot the kymographs of the simulated cell at different values of activity parameter $\beta_0$ with large noise parameter $\sigma_v$, as shown in Fig.4 in the absence of noise. We find that the larger noise does not alter the basic characteristics of the different migration modes on the adhesion gradient.

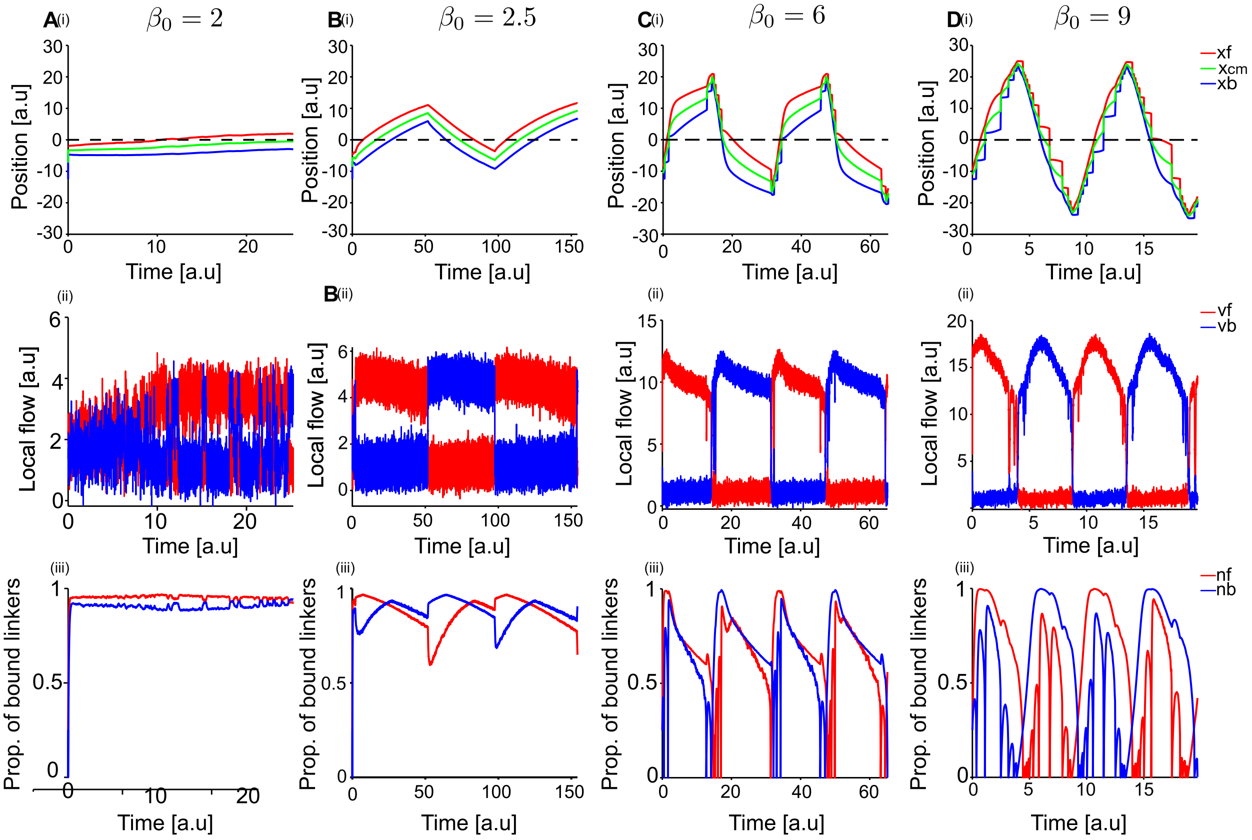


Figure S1. **Four modes of migration on exponential adhesion gradient ($\sigma_v = 0.4$).** **A**[(i)-(iii)] Time series plots at $\beta_0 = 2$ showing the cell trapped at the peak. (i) Position time series: front (red), center of mass (green), and back (blue) of the cell; dotted line indicates the peak of the adhesion gradient. (ii) Local actin flow at the front (red) and back (blue). (iii) Number of bound linkers at the front (red) and back (blue). **B**[(i)-(iii)] Time series plots at $\beta_0 = 2.5$ showing smooth oscillatory migration around the peak. **C**[(i)-(iii)] Time series plots at $\beta_0 = 6$ showing stick-slip migration while going up the gradient and smooth migration while moving down the gradient. **D**[(i)-(iii)] Time series plots at $\beta_0 = 7$ showing only stick-slip migration. Model parameters: $A = 27$, $k_g = 0.1769$, $\tilde{\alpha} = 1$, $r_0 = 1$, $\sigma_v = 0.4$.

### S2. PHASE SPACE DIAGRAM AT HIGHER NOISE LEVEL ($\sigma_v = 0.4$)

In Fig.S2 we show the effects of higher noise on the trajectory of the cell along an exponential adhesion gradient, as was shown in Fig.5 in the absence of noise. The larger noise makes the stick-slip migration exhibit larger and fewer jumps up the adhesion gradient, but qualitatively the behavior remains the same as in the absence of noise.

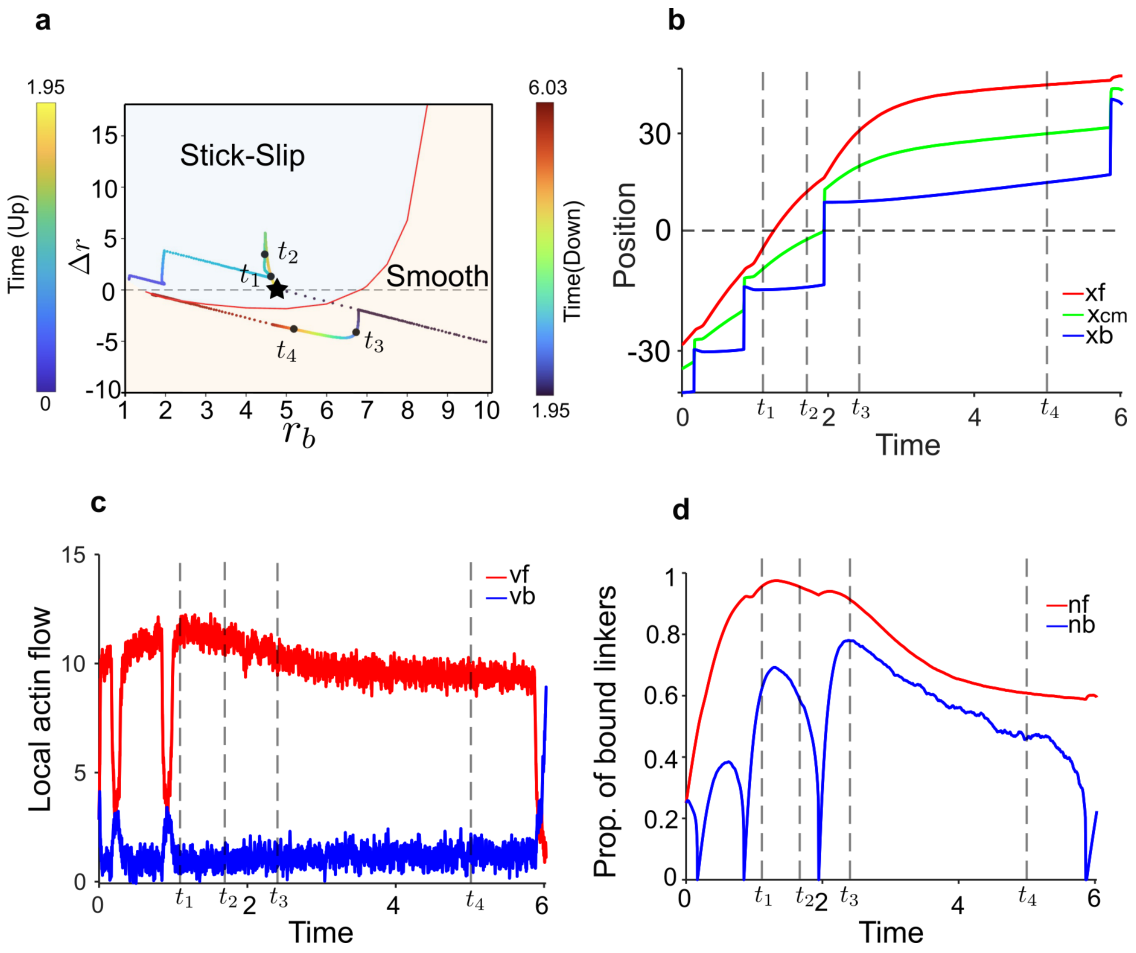


Figure S2. **Phase space diagram at $\beta_0 = 6$ ($\sigma_v = 0.4$).** **a** $\Delta r$ versus $r_b$ phase space diagram, where time series trajectories are shown as a color gradient from blue ($t = 0$) to yellow ($t = 1.95$) as the cell moves up the gradient, and from blue ($t = 1.95$) to red ($t = 6.03$) as the cell moves down the gradient. Stick slip and smooth regimes are marked in steel blue and cream colors respectively and the transition line is marked in red. Times $t_1 = 1.11$, $t_2 = 1.7$, $t_3 = 2.4$ and $t_4 = 5$ are marked in both the phase space diagram and subsequent time series plots, showing stick-slip mode at $t_1$ and $t_2$, and smooth mode at $t_3$ and $t_4$. The point at which the cell crosses the peak is marked with a black star. **b** Position time series plot with marked times shown as vertical dotted lines. Positions of the front, center, and back are shown in red, green, and blue, respectively. **c** Local actin flow time series plot with marked time points shown as vertical dotted lines. Flow at the front and back are shown in red and blue, respectively. **d** Fraction of bound linkers time series plot with marked time points shown as vertical dotted lines. Front and back values are shown in red and blue, respectively. Model parameters: $\beta_0 = 6$, $A = 10$, $k_g = 0.1769$, $\tilde{\alpha} = 1$, $r_0 = 1$, $\sigma_v = 0.4$.

### S3. HAPTOTAXIS AND ANTI HAPTOTAXIS PERCENTAGES AT DIFFERENT $\beta_0$ VALUES

In Fig.S3 we show the dependence of the haptotactic bias on the activity ($\beta_0$) and the noise ($\sigma_v$) parameters of the cell.

For low activity ($\beta_0 = 2$ and $\beta_0 = 2.5$), the cells move slowly and lack strong active polarization, instead they end up drifting up the adhesion gradient. Because their haptotactic bias does not depend on maintaining polarization, noise has little additional disruptive effect therefore haptotactic percentages remains high and and stable ( 80-95%) across all noise levels.

At intermediate activity ($\beta_0 = 4$), cells develop weak polarization, placing them in vulnerable regime where even moderate noise can override their directional bias, causing a noticeable dip in the haptotactic percentages with increasing noise magnitude. Interestingly, at very high noise the haptotactic bias recovers, because strong fluctuations destroy polarization entirely and cells revert to passive gradient-driven drift.

At higher activity ($\beta_0 = 6.5, 9$ and 11), cells are strongly polarized and commit to migration in a single direction. While the strong polarization provides some resistance to noise, it also means that when the noise is sufficient to flip the committed migration direction, the effect is more consequential as cells become polarized persistently in the anti-haptotactic direction. Consequently, haptotactic percentages start near 100% at low noise but decline more steeply before stabilizing around $55\% - 75\%$ at high noise, without the recovery seen at the intermediate $\beta_0$. At intermediate noise for higher $\beta_0$ we find that the region of weak directionality increases over a wider range of noise level. This further demonstrates the stochastic nature of these direction-switching events. Therefore these results suggest that gradient sensing is not simply a function of how active (and persistent) a cell is but rather emerges from an interplay between polarization strength and susceptibility to noise.

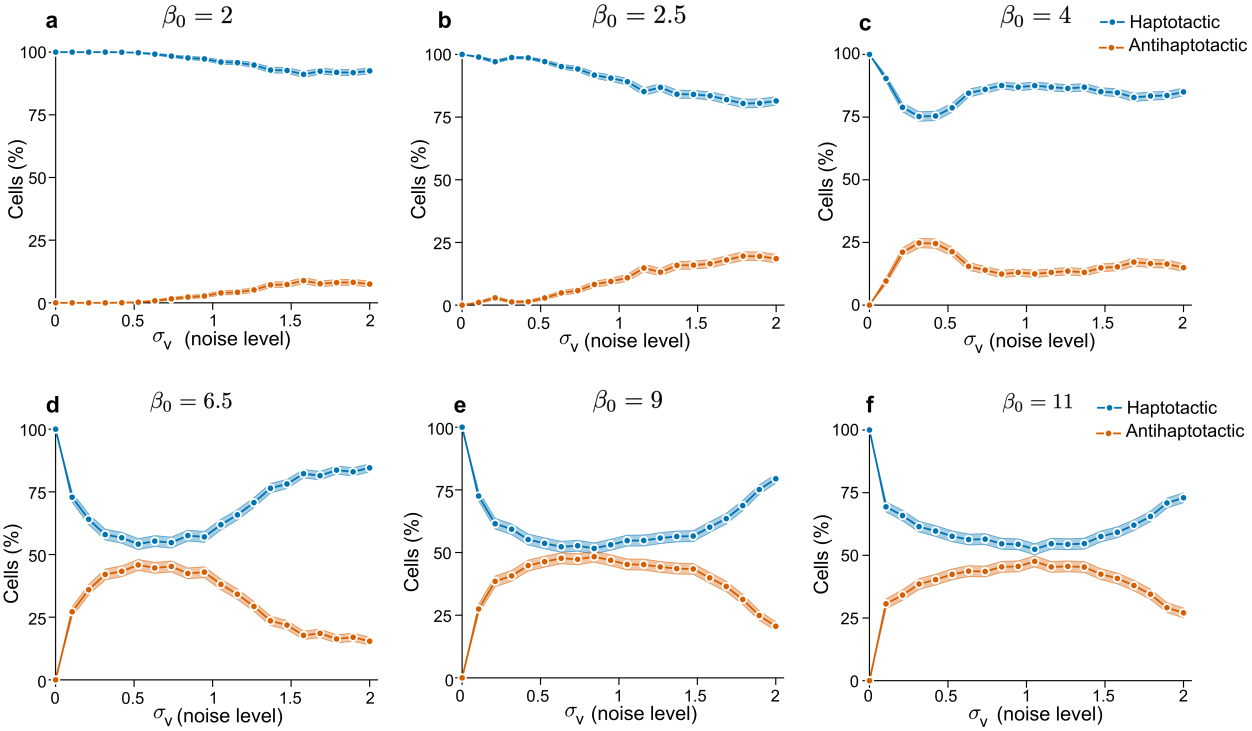


Figure S3. **Noise dependence of directionality across baseline activity levels ($\beta_0$).** Haptotactic (blue) and anti-haptotactic (orange) fractions are shown as a function of the velocity-noise amplitude $\sigma_v$ for different baseline polymerization activities $\beta_0$ (panels a–f). Shaded bands denote variability across stochastic realizations (computed over repeated simulations at fixed parameters). Model parameters: $A = 27$, $k_g = 0.1769$, $\tilde{\alpha} = 1$, $r_0 = 1$, track length $= 65$. Statistics are computed from N $=$ 2000 simulated trajectories.

### S4. EFFECT OF ADHESION-GRADIENT MAGNITUDE ON CELL LENGTH

In Fig.S4 we show the effect of increasing the magnitude of the adhesion maximum (and therefore of the adhesion gradient), on the mean cell length. The adhesion levels correspond to those shown in Fig.9. We find, as expected, that on the more adhesive substrates cells tend to have higher spreading and elongation.

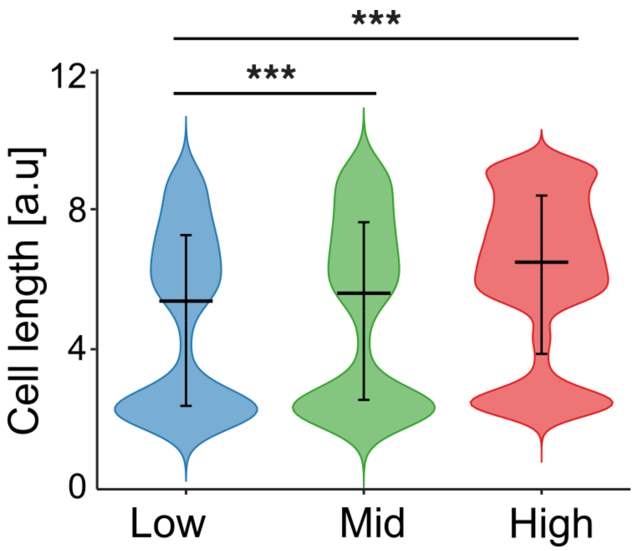


Figure S4. **Effect of adhesion-gradient magnitude on model cell length distributions.** Error bars represent standard deviation, significant differences were identified using the Mann–Whitney $U$ test (***$P < 0.001$). Model parameters: Low, mid and high adhesion magnitude values are: $A = 22, 27, 32$ respectively, $k_g = 0.1769$, $\beta_0 = 7.5$, $\tilde{\alpha} = 1$, $r_0 = 1$, $\sigma_v = 0.4$. Statistics are computed from $N = 2000$ simulated trajectories.

### S5. TRACTION FORCES

A traction kymograph maps the spatiotemporal distribution of traction forces exerted by the cell on the substrate, which can be measured experimentally using traction force microscopy [1, 2]. In the model, traction forces are evaluated at the front and rear adhesion sites. For more realistic plotting, these forces are distributed over a spatial domain by convolution with Gaussian functions. The resulting kymograph displays these forces as a heatmap with the cell-boundary trajectories overlaid.

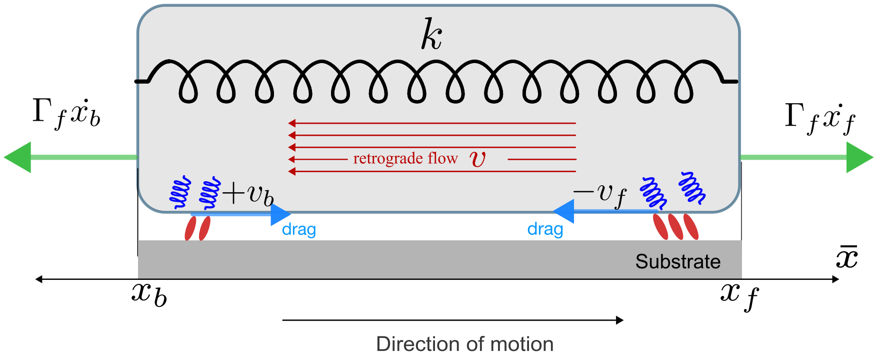


Figure S5. **Schematic representation of traction-force calculation.** Traction forces are evaluated at the front and rear adhesion sites and distributed spatially within the cell body using Gaussian profiles.

From the model equations (Eqs.1-5), the traction forces at the front and rear boundaries are written as

$$T_f = -\frac{r_f}{r_f + r_0} v_f + \Gamma_f \dot{x}_f, \tag{S1}$$

$$T_b = +\frac{r_b}{r_b + r_0} v_b + \Gamma_b \dot{x}_b. \tag{S2}$$

Here, $r_f$ and $r_b$ are the adhesion strengths at the front and rear, respectively, $v_f$ and $v_b$ are the local actin flows, and $\dot{x}_f$ and $\dot{x}_b$ are the velocities of the front and rear cell boundaries. The clutch-dependent friction coefficients $\Gamma_f$ and $\Gamma_b$, defined in Eq.5, determine how efficiently the actin flow is transmitted to the substrate.

To construct a continuous traction field, the boundary forces are distributed using normalized Gaussian functions $\mathcal{N}_f(x)$ and $\mathcal{N}_b(x)$, centered near the front and rear adhesion sites:

$$T(x, t) = T_f \mathcal{N}_f(x) + T_b \mathcal{N}_b(x). \tag{S3}$$

The Gaussian width is set proportional to the cell length,

$$w = \frac{x_f - x_b}{8},$$

with centers positioned at

$$\mu_f = x_f - 2.5w, \qquad \mu_b = x_b + 2.5w,$$

so that the traction forces are distributed within the cell body rather than exactly at the cell edges.

A few examples are shown in Fig.15 in comparison with experiments. In Fig.S6 the traction force kymographs are shown for the main migration modes, corresponding to Fig.4.


[1] P. Roca-Cusachs, V. Conte, and X. Trepat, Quantifying forces in cell biology, Nature cell biology **19**, 742 (2017).

[2] M. Dembo and Y.-L. Wang, Stresses at the cell-to-substrate interface during locomotion of fibroblasts, Biophysical journal **76**, 2307 (1999).

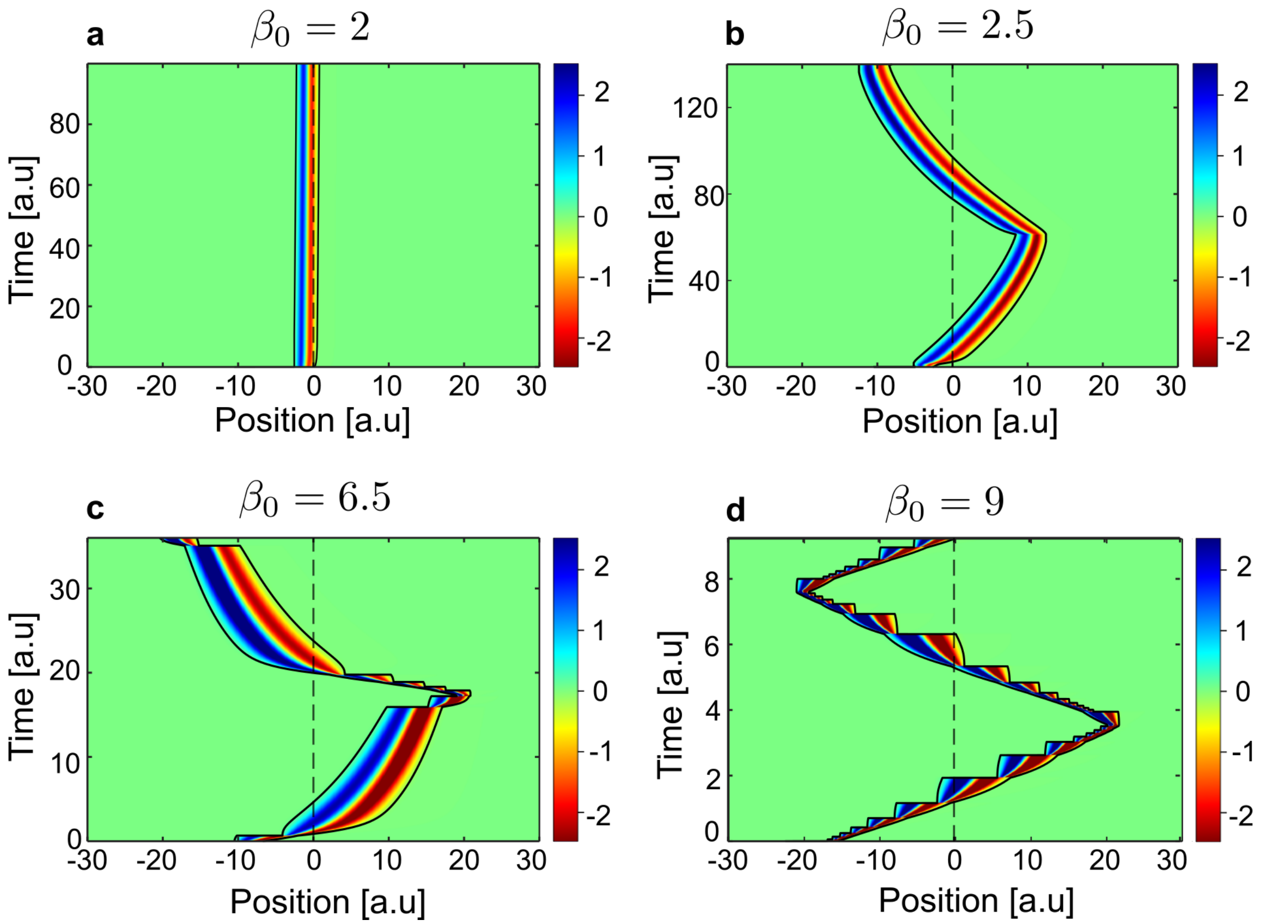


Figure S6. **Traction kymograph for the four main modes of migration.** **a**) $\beta_0 = 2$: the cell remains stuck at the peak adhesion position, marked by the dotted line. **b**) $\beta_0 = 2.5$: the cell shows smooth oscillatory migration around the peak. **c**) $\beta_0 = 6.5$: the cell exhibits stick-slip migration while moving up the adhesion gradient and smooth migration while moving down. **d**) $\beta_0 = 9$: the cell shows stick-slip migration in both directions. Model parameters: $A = 27$, $k_g = 0.1769$, $\tilde{\alpha} = 1$, $r_0 = 1$, $\sigma_v = 0.01$. Peak adhesion position: $x_{\text{peak}} = 0$, cell track size $= 65$. Red and blue indicate traction forces of opposite sign, while green indicates approximately zero traction.
